# G1 Phase Lengthening During Neural Tissue Development Involves CDC25B Induced G1 Heterogeneity

**DOI:** 10.1101/2020.11.06.370833

**Authors:** Angie Molina, Frédéric Bonnet, V. Lobjois, Sophie Bel-Vialar, Jacques Gautrais, Fabienne Pituello, Eric Agius

## Abstract

While lengthening of the cell cycle and G1 phase is a generic feature of tissue maturation during development, the underlying mechanism remains still poorly understood. Here we develop a time lapse imaging strategy to measure the four phases of the cell cycle in single neural progenitor cells in their endogenous environment. Our results show that neural progenitors possess a great heterogeneity of the cell cycle length. This duration variability is distributed over all phases of the cell cycle, with the G1 phase being the one contributing primarily to cell cycle variability. Within one cell cycle, each phase duration appears stochastic and independent except for a surprising correlation between S and M phase. Lineage analysis indicates that the majority of daughter cells display longer G1 phase than their mother’s suggesting that at each cell cycle a mechanism lengthens the G1 phase. We identify an actor of the core cell cycle machinery, the CDC25B phosphatase known to regulate G2/M transition, as an indirect regulator of the duration of the G1 phase. We propose that CDC25B acts via a cell to cell increase in G1 phase heterogeneity revealing a novel mechanism of G1 lengthening associated with tissue development.

## INTRODUCTION

Building a multicellular functional organ requires tight coordination between cell proliferation, cell fate specification and differentiation. In the developing nervous system, the spatio-temporal regulation of these processes is of key importance to generate the relative proportion of the numerous classes of neurons and glia cells essential to construct functional neuronal circuits.

The cell cycle and components of the core cell cycle machinery have been shown to play a major role in the decision to proliferate or differentiate in embryonic stem cells, pluripotent stem cells and neural stem/progenitor cells (for a review see Liu, et al., 2019). In numerous cell types including neural stem/progenitor cells, cell cycle and G1 phase lengthening is a general feature accompanying cell maturation and differentiation. During mammalian corticogenesis where consecutive types of progenitors have been described, lengthening of the G1-phase is associated with the transition from neural-stem-like apical progenitors (AP) to fate restricted basal progenitors (BP) and a shortening of the S phase with the transition from proliferative to neurogenic divisions (Arai, et al., 2011). Reducing G1 phase length results in an inhibition of neurogenesis, while lengthening G1 duration promotes neurogenesis (Lim and Kaldis, 2012; Artegiani, et al., 2011; Lange, et al., 2009; Pilaz, et al., 2009). In the developing spinal cord, different cell cycle kinetics are observed in discrete domains of neural progenitors (Molina and Pituello, 2017). Differentiation here progresses from ventral to dorsal with time. When the maximum differentiation rate is reached in the ventral domain, neural progenitor cells (NPCs) exhibit a long G1 phase and short S and G2 phases (Kicheva, et al., 2014; Saade, et al., 2013; Peco, et al., 2012). In contrast, the dorsal domain mainly composed at the same age of proliferative NPCs, shows a short G1 phase accompanied by long S and G2 phases (Kicheva, et al., 2014; Saade, et al., 2013; Peco, et al., 2012). Overexpressing D-type Cyclins in young neural tube increases the pool of proliferating progenitors and induces a transient reduction of neuron production (Lacomme, et al., 2012; Cao, et al., 2008), while more mature NPCs will differentiate regardless of Cyclin D overexpression and cell cycle exit (Lobjois, et al., 2008). Shortening of the G2 phase associated with neurogenesis results from the upregulation in NPCs of a regulator of the G2/M transition, the CDC25B phosphatase. CDC25B promotes entry into mitosis by dephosphorylating its canonical substrates, the cyclin-dependent kinases (CDKs) complexes. Surprisingly for a positive regulator of the core cell cycle machinery, CDC25B has been shown to promote neurogenesis in mouse, chicken and Xenopus embryos (Peco, et al., 2012; Gruber, et al., 2011; Ueno, et al., 2008). Gain and loss of function experiments performed in chicken neural tube show that CDC25B induces the conversion of proliferating NPCs into differentiating neurons by promoting neurogenic divisions (Bonnet, et al., 2018). CDC25B acts using both CDK independent and dependent molecular mechanisms (Bonnet, et al., 2018). A mathematical model has allowed us to hypothesize that CDC25B expression in neural progenitors progressively restricts the proliferative capacities of embryonic neural stem cells (Azaïs et al., 2019).

A weak point common to all these studies is that cell cycle analyses have been performed at the population level. Cell cycle and phases lengths were calculated from fixed tissues and correspond to estimated average length values considering the population of NPCs as homogeneous (Nowakowski, et al., 1989). Whether this neural progenitor population is homogeneous or heterogeneous relative to cell cycle kinetics is poorly documented. An indication that the population of NPCs is indeed heterogeneous comes from a study where the total cell cycle length (Tc) was determined from the length of time measured between two cell divisions using time-lapse imaging in chick embryo slice cultures (Wilcock, et al., 2007). These single cell measurements reveal that the neural progenitor population displays marked heterogeneity regarding the Tc, which ranges from 9 hours to 28 hours. Moreover, previous data indicate that the cell cycle length of cells dividing to produce two progenitors was shorter than that producing one neuron and one progenitor (Wilcock, et al., 2007).

The goal of this study was to determine at the cellular level the features of NPCs cell cycle kinetics in their endogenous environment and the link between cell cycle kinetics and cell aging. To do so, we set up a high-resolution time-lapse imaging technique to measure the duration of each phase of the cell cycle in single NPCs within the developing neural tube and to track the behavior of daughter cells after mitosis. We found that the NPCs population is highly heterogeneous regarding the distribution of cycling time. We showed that the duration of each phase can be described by a stochastic process and found no coupling between phase length within a cell cycle. All phases, but M phase, are very heterogeneous, and the total cell cycle (Tc) length variation can be primarily explained by G1 heterogeneity. Time lapse imaging showed that the majority of daughter cells display a longer G1 phase than their mother’s suggesting that gradually at each cell cycle a mechanism lengthens the G1 phase. Because CDC25B affects both cell cycle kinetics and neurogenesis, we tested its effect and showed that it increases cell-to-cell G1 length heterogeneity, thereby raising tissue-scale G1 length without affecting cell phase independence. We propose that in a context of cell cycle phases independence, CDC25B expression and reiteration at each cell cycle, increases heterogeneity in G1 phase durations and promotes the restriction of the proliferative capacities of embryonic neural stem cells, their maturation and commitment to differentiation.

## MATERIALS AND METHODS

### Embryos

Fertile hens’ eggs were incubated at 38°C in a humidified incubator to yield embryos appropriately staged (Hamburger and Hamilton, 1992).

### DNA constructs and *in ovo* electroporation

*In ovo* electroporation experiments were performed using 1.5- to 2-day-old chicken as described previously (Peco, et al., 2012). To detect the four phases of the cell cycle we developed the mKO-zCdt1-pIRES-NLS-eGFP-L2-PCNA biosensor. In the pCAG plasmid, we inserted the Fucci G1 probe derived from zebrafish Cdt1 (Sugiyama, et al., 2009) and downstream the IRES element, the NLS green fluorescently tagged proliferating cell nuclear antigen (PCNA, Leonhardt, et al., 2000). This FUCCI G1-PCNA vector is transfected at 0.5μg/μl by *in ovo* electroporation in the chicken neural tube to reproducibly obtain a high degree of mosaicism compatible with lineage tracing (Wilcock, et al., 2007). The hCDC25B or hCDC25B^ΔCDK^ gain-of-function experiments were performed at 1.5μg/μl as in Bonnet, et al., 2018. The Fucci S/G2/M mAG-hGem (Sakaue-Sawano, et al., 2008) and the fucci G1 mKO2-zCdt1 (1/190) (Sugiyama, et al., 2009) were used at 0.5 μg/μl.

### Immunohistochemistry

Embryos or neural tube explants are fixed in 3.7% formaldehyde for 2 hours and sliced using a vibratome (Leica). Proteins were detected on 50 μm vibratome sections, as previously described (Peco, et al., 2012). The antibodies used are: anti-Olig2 (Milipore), anti-Islet1/2, anti-MNR2 (Developmental Studies Hybridoma Bank), anti-active caspase 3 (Becton Dickinson Biosciences).

Determination of S-phase length (Ts) and total cell cycle length (Tc) were based on the relative numbers of cells that incorporated one or two thymidine analogs (Martynoga et al., 2005). For in ovo incorporation 10 μl of bromodeoxyuridine (BrdU Sigma, St Louis, MO, USA) (500 μM) were injected into embryos and followed after 90 minutes by EdU (500 μM, Invitrogen) incorporation. Embryos were fixed 30 minutes later. EdU was detected first (Click-iT EdU Alexa Fluor 647 Imaging Kit, Invitrogen), followed by BrdU detection using the G3G4 antibody that does not recognize EdU. For BrdU incorporation, 10 μl of BrdU (500 μM) were injected into embryos, reincubated for 30 minutes before fixation. BrdU immunodetection was performed on vibratome sections using anti-BrdU (mouse monoclonal, G3G4) as in (Lobjois, et al., 2004). Cell death was analyzed by immunofluorescence, using the anti-active caspase 3 antibody.

### Flow cytometry analysis

1.5- to 2-day-old chicken embryos were electroporated with H2B-GFP or NLS-eGFP-L2-PCNA constructs. Neural tubes were dissected 24 h following electroporation, incubated at 37°C for 10 min in trypsin-EDTA to obtain a single-cell suspension, and fixed for 30 minutes in 4% formaldehyde. Cell suspensions were incubated for 30 min in 400 μl of propidium iodure (PI) (20μg/ml)/ RNAse (100μg/ml) cocktail (Sigma). PI and GFP fluorescence were acquired with a FACSCalibur cytometer (Cat#342975, Becton Dickinson), and DNA content analysis was performed using the FlowJo software.

### Embryonic neural tube culture and Time-lapse Imaging

Electroporated E2 embryos were collected in PBS and 100 μm slices were obtained using a McIlwain tissue chopper (WPI), from the brachial region corresponding to somites 12 to 17 which generates the greatest number of motoneurons (Oppenheim, et al., 1989). Sections were collected in 199 culture medium (GIBCO) and were sorted out under a fluorescence microscope to control tissue integrity and the presence of isolated fluorescent cells along the dorso-ventral axis. Each slice was embedded into 10 μl of rat type I collagen (Roche Diagnosis; diluted at 80% with 1X MEM (GIBCO), 1X GlutaMax (GIBCO) and neutralizing bicarbonate (GIBCO). 4 neural tube-containing collagen drops (5 μl) were distributed on a 35 mm glass-bottom culture dish (IBIDI) (procedure modified from Das, et al., 2012. Collagen polymerization was performed at 38°C for 30 minutes and 1.5 ml of complete culture medium is added (Medium 199 (Gibco) supplemented with 1X GlutaMax (Gibco), 5% Fetal calf Serum (FCS, Fisher Scientific) and Gentamycin (Gibco, 40ug/mL). The culture dish is placed in a humid atmosphere incubator with 5% CO2 for 12 hours before time lapse imaging. Alternatively, explants were cultured for 24 or 48 hours and then fixed in 3.7% formaldehyde (FA) and processed for immunostaining.

For time-lapse, images were acquired on an inverted microscope (Leica inverted DMI8) equipped with a heating enclosure (set up at 39°C) in an atmosphere containing 5% CO2, a spinning disk confocal head (CSU-X1-M1N, Yokogawa), a SCMOS camera and a 63X oil immersion objective (NA 1,4 −0,7). Attenuation of laser beam pulses were performed to reduced cell damage due to photo-toxicity (Boudreau, et al., 2016). We recorded 40 μm thick z stacks (2 μm z-steps) at 5 min intervals for 48h.

### Imaging, data analysis and statistics

IMARIS® and ImageJ® softwares were used for image processing and data analysis. Statistical analyses were performed using GraphPad Prism and R software. Details for methods used in SI-Data are given in the relevant sections. The normality of the data sets was determined, analyses of variance performed and the appropriate t-test method used. Values shown are mean ± standard error of the mean (s.e.m.). Significance was assessed by performing the Student-Mann-Whitney test. The significance values are: *P<0.05; **P<0.01; ***P<0.001. See also SI.

## RESULTS

### A new method to measure the four cell cycle phases length of NPCs in real time

In order to determine cell cycle kinetics of individual spinal NPCs, we developed a combination of biosensors to detect unambiguously the four phases of the cell cycle in living cells. To label the G1 phase, we used the zebrafish G1 marker mKO2-zCdt1 (FUCCI G1, (Sugiyama, et al., 2009) instead of the human Cdt1, which in chick NPCs, persists in all cell cycle phases, suggesting that it is not properly degraded (data not shown). We verified the zebrafish FUCCI G1 specificity in G1 by co-electroporating mKO2-zCdt1 with FUCCI S/G2/M (mAG-hGeminin) and quantifying, 24 hours later, red, green and yellow cells corresponding to cells expressing respectively FUCCI G1, FUCCI S/G2/M or both (Fig. 1A). We quantified 43,7% and 41,4 % of cells expressing FUCCI G1 or FUCCI S/G2/M respectively and 14,9 % expressing both markers illustrating their transient overlap as already reported (Sakaue-Sawano, et al., 2008). This result was comparable to previous quantifications performed by flow cytometry analysis (FACS, Benazeraf, et al., 2006). To time more precisely the G1/S transition and to identify the S/G2 transition, we used NLS-eGFP-L2-PCNA protein (Leonhardt, et al., 2000). S phase onset was detected by the punctuated appearance of NLS-eGFP-L2-PCNA. PCNA is recruited within the DNA replication forks that change in location and size as S phase progresses. A 30 minute BrdU pulse confirmed that the punctate labelling observed with NLS-eGFP-L2-PCNA corresponds to S phase cells (Fig. 1B). We tested whether electroporated NLS-eGFP-L2-PCNA induces cell cycle disturbances by means of FACS analysis using propidium iodide as a DNA intercalating agent. As shown in Fig. 1C, there is no difference in the cell cycle profile or in the corresponding quantifications following NLS-eGFP-L2-PCNA or H2B-GFP electroporation. These results confirm that NLS-eGFP-L2-PCNA overexpression does not interfere with cell cycle timing, as previously reported in zebrafish neuroepithelia (Leung, et al., 2011). According to this, we constructed a pCAG plasmid containing mKO-zCdt1 and NLS-eGFP-L2-PCNA (Fig. 1D) to perform proper mosaic expression through *in ovo* electroporation into 2-day-old chick neural tube (Fig.1E), and thereby visualize the four cell cycle phases in individual cycling neural progenitors (Fig.1F).

**Figure 1.**
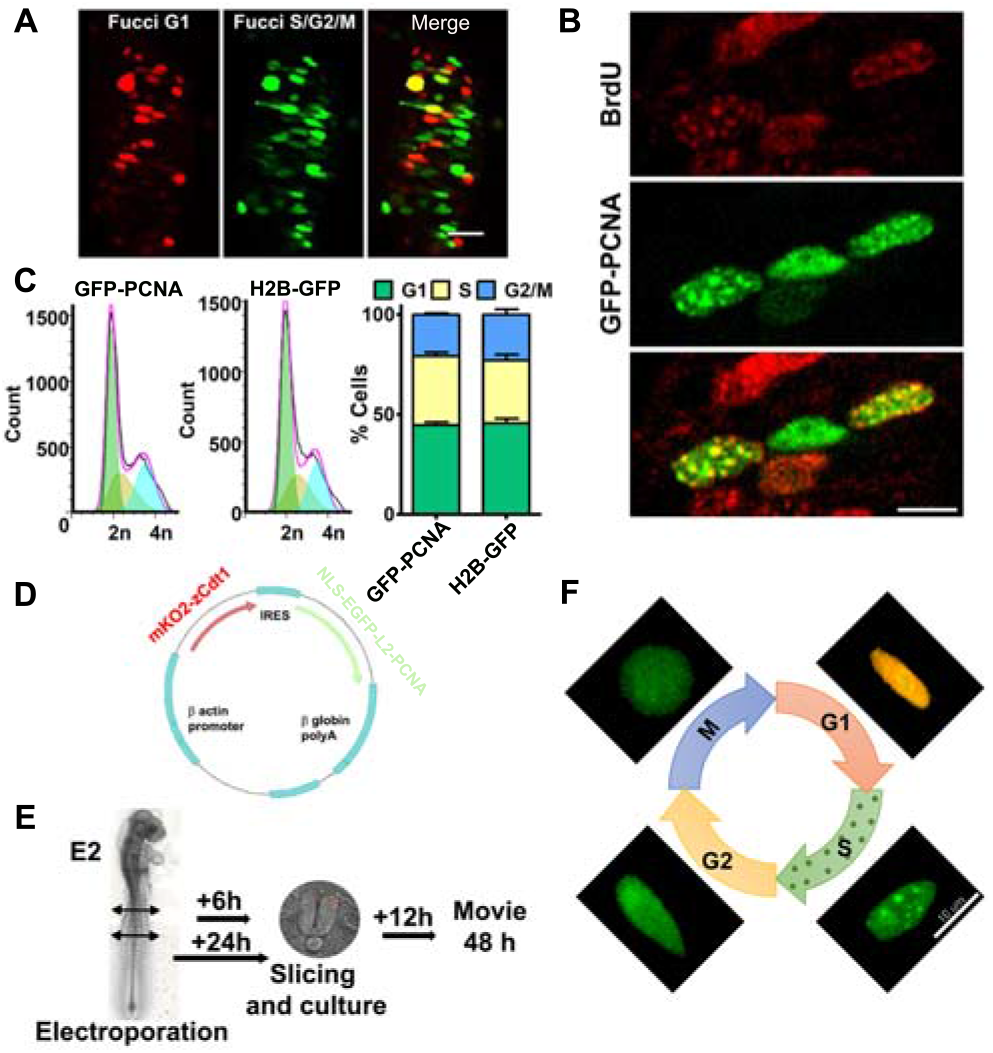
FUCCI-G1-PCNA biosensor marks the four cell cycle phases in chick neuroepithelium. **A)** Cross-sections of chick neural tube expressing FUCCI G1 and FUCCI S/G2/M vectors. **B)** Representative sections of chick embryonic spinal cord expressing GFP-PCNA and stained after BrdU incorporation to identify S-phase cells. **C)** FACS profiles of DNA content for GFP-PCNA or H2B-GFP electroporated NPCs (left panel). DNA histograms made from PI staining and FACS flow cytometry (right panel). Percent cells in G1 are 44.7 ± 1.4% compared to 45.7 ± 2.4%. Percent cells in S are 34.5 ± 1.9% compared to 31.4 ± 2.9% and Percent cells in G2/M phases are 20.8 ± 0.7% compared to 22.9 ± 2.6% in GFP-PCNA and H2B-GFP respectively. **D)** Schematic representation of the mKO-zCdt1-pIRES-NLS-EGFP-L2-PCNA biosensor. **E)** Scheme of the experimental protocol for time-lapse imaging. **F)** Representative images of chick NPC nuclei through the cell cycle after electroporation of FUCCI G1-PCNA vector. The four phases are identified by differential expression and distribution of FUCCI-G1 and GFP-PCNA proteins. Scale bar = 10 μm.

To analyze cell cycle parameters, E2 embryonic neural tubes were electroporated with the FUCCI G1-PCNA vector. After 6 hours (E2.25) or 24 hours (E3) of incubation, embryos were sliced, and tissue cultures recovered for 12 hours prior imaging (Fig. 1E). This step of recovering is important to bypass the lengthening of the cell cycle and the delayed increase in the population of progenitors and neurons observed following slicing and cultivation (Sup Fig1 A-D). Time-lapse imaging was performed for 48 hours using a confocal microscope. It is important to stress that the use of 5% fetal calf serum in the culture medium and using spinning disk confocal microscopy associated with attenuation of laser beam pulses to reduce phototoxicity (Boudreau, et al., 2016) are critical to perform cell cycle measurements. Visualization of the FUCCI G1-PCNA vector by time-lapse imaging revealed that the different transitions in the cell cycle could be discerned using nuclear expression of both proteins (Fig. 1F): G1 phase was characterized by the co-expression of mKO-FUCCI G1 and NLS-eGFP-L2-PCNA. S phase was detected by the appearance of punctuated NLS-eGFP-L2-PCNA associated with the gradual disappearance of the FUCCI-G1 reporter. In G2 phase, NLS-eGFP-L2-PCNA was evenly distributed inside the nuclei, and finally mitosis was detected by nuclear envelope breakdown accompanied by morphology changes of NPCs, which become rounded (Fig. 1F).

We have thus designed a long-term time-lapse imaging method to measure accurately the four cell cycle phases in individual cycling neural progenitors in an endogenous environment, the neural tube.

### Neural progenitor nuclei display INMs in phase with the cell cycle and three distinct behaviors after mitosis

Electroporation and imaging of the FUCCI G1-PCNA vector in E2 (HH12) stage embryos, showed cycling NPCs displaying G1 nuclei (orange) moving to the basal side, nuclei in S phase (green punctuated) located in the basal half of the ventricular zone and G2 nuclei (green) moving back to the apical side where mitosis occurs (Fig. 2A; Movie 1). These characteristic positions indicate that in neuroepithelia, INM occurs in phase with the cell cycle (Fig. 2B) as previously described by a large number of studies (Molina and Pituello, 2017; Laguesse, et al., 2015; Kosodo, et al., 2011; Langman, et al., 1966). We also observed nuclei on the basal side of the neural tube expressing brighter orange fluorescence (Fig. 2A), probably due to the accumulation of FUCCI G1 in differentiating G0 neurons after cell cycle exit, as previously described (Sakaue-Sawano, et al., 2008).

**Figure 2.**
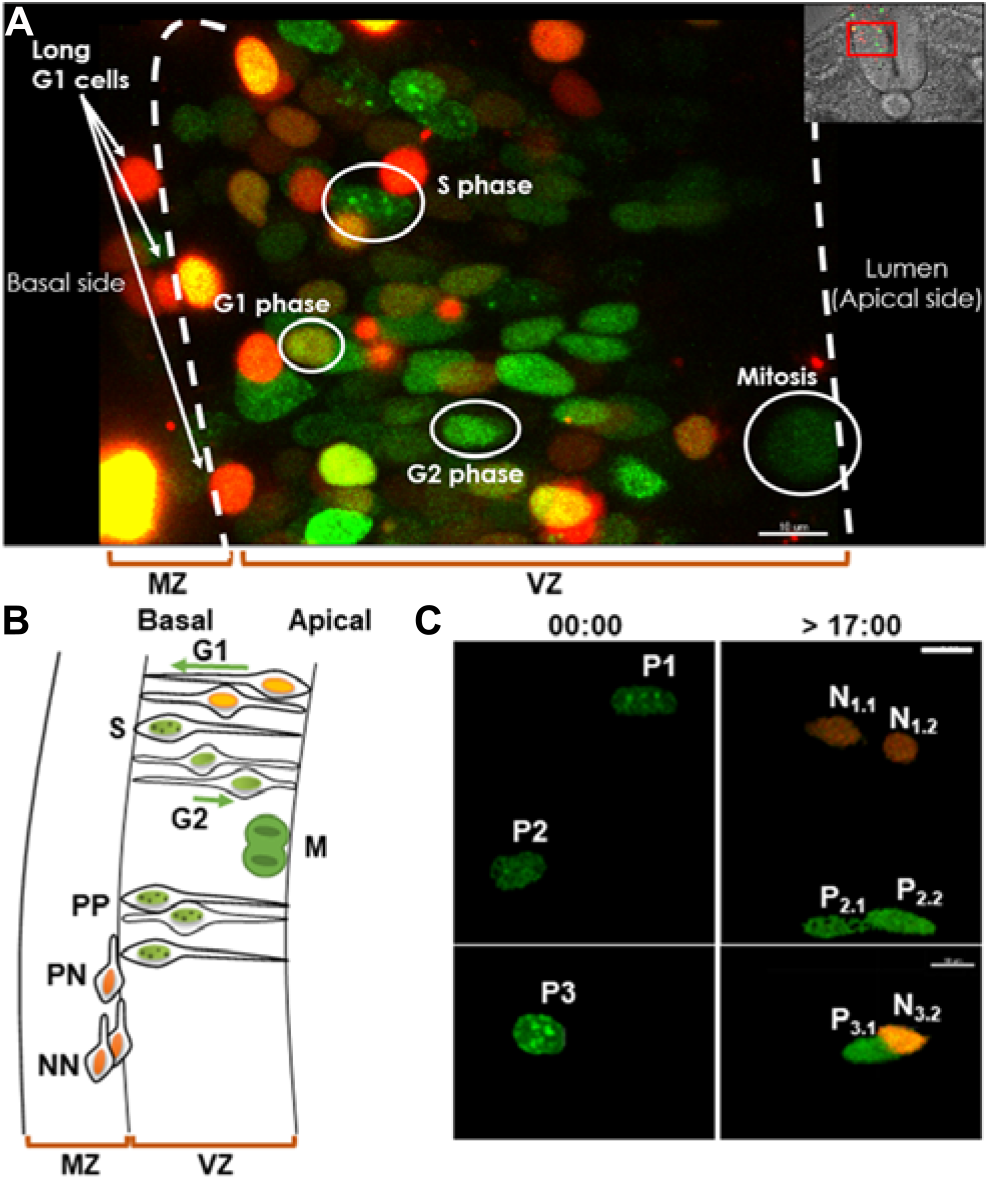
Time lapse observation of NPCs displaying three types of cell division after mitosis. **A)** Still picture of a time lapse video showing the expression of the sensor in the nucleus of NPCs that allows identification of the four cell cycle phases and the corresponding position within the neuroepithelium. **B)** Schematic representation of the interkinetic nuclear movement (INM) and of the three modes of division (PP, PN and NN) occurring in the neural tube. MZ, mantle zone. VZ, ventricular zone. **C)** Still images of a E2.25 culture time lapse movie showing the three modes of divisions observed in the spinal cord from progenitors P1, P2, P3: proliferative (PP - P_2.1_ - P_2.2_), asymmetric neurogenic (PN - P_3.1_ - N_3.2_) and terminal neurogenic (NN - N_1.1_ - N_1.2_). Scale bars represent 10 μm.

Nuclear tracking after mitosis reveals three different behaviors in daughter cells (Fig. 2C): both daughter cells re-enter S phase and nuclei switch from FUCCI G1 to PCNA-punctuated labeling, this behavior clearly corresponds to cells performing proliferative divisions (PP) (P2 giving P2.1 and P2.2); only one daughter cell re-enters the cell cycle, while the nucleus of the other remains orange and will migrate to the periphery (P3 giving P3.1 and N3.2); the two daughter cells nuclei remain orange and will migrate to the periphery (P1 giving N1.1 and N1.2). These orange nuclei located at the basal side and displaying a G1 phase longer than 1000 min (16h40), were never observed re-entering S phase. We thus assumed that they correspond to cells primed to differentiate into neurons (Fig. 2A-Movie 1). The G1 lengths measured for these cells are excluded from our analyses. Hence our strategy allows us to follow the nuclei of NPCs across the cell cycle and to determine, after mitosis, whether the daughter cells re-enter the cell cycle or remain in G1 phase for a long time (marked PL_G1_).

### Cell cycle kinetics are highly heterogeneous in the population of NPCs

Cell cycle kinetics analysis was performed with E2 (HH12) stage embryo to ensure that we analyze young proliferative progenitors (Fig.3A). We first compared, at the population level, the cell cycle data measured using time-lapse imaging with those obtained on fixed tissues. The mean value measured using time lapse for the total cell cycle (Tc) is 14h01 (n=33), G1: 5h09 (n=50), G2: 1h17 (n=54) and M : 31 min (n=50, Table 1). These data are in agreement with those described for fixed embryos at equivalent developmental stages (brackets in Fig. 3B), Tc : 10h and 16h, G1 : 4h30 and 7h, G2 : 1h18 min and 2h and mitosis : 30 minutes (Kicheva, et al., 2014; Le Dreau, et al., 2014; Saade, et al., 2013; Peco, et al., 2012; Wilcock, et al., 2007). The S phase length measured using time lapse (7h18 ± 23 min) presents an average value higher than reported in fixed tissues (3h42 min and 5h54 min). Thus, except for this phase, which was slightly increased in our conditions (see discussion), the mean data obtained at the population level are consistent with those reported for fixed tissues at equivalent developmental stage.

**Figure 3.**
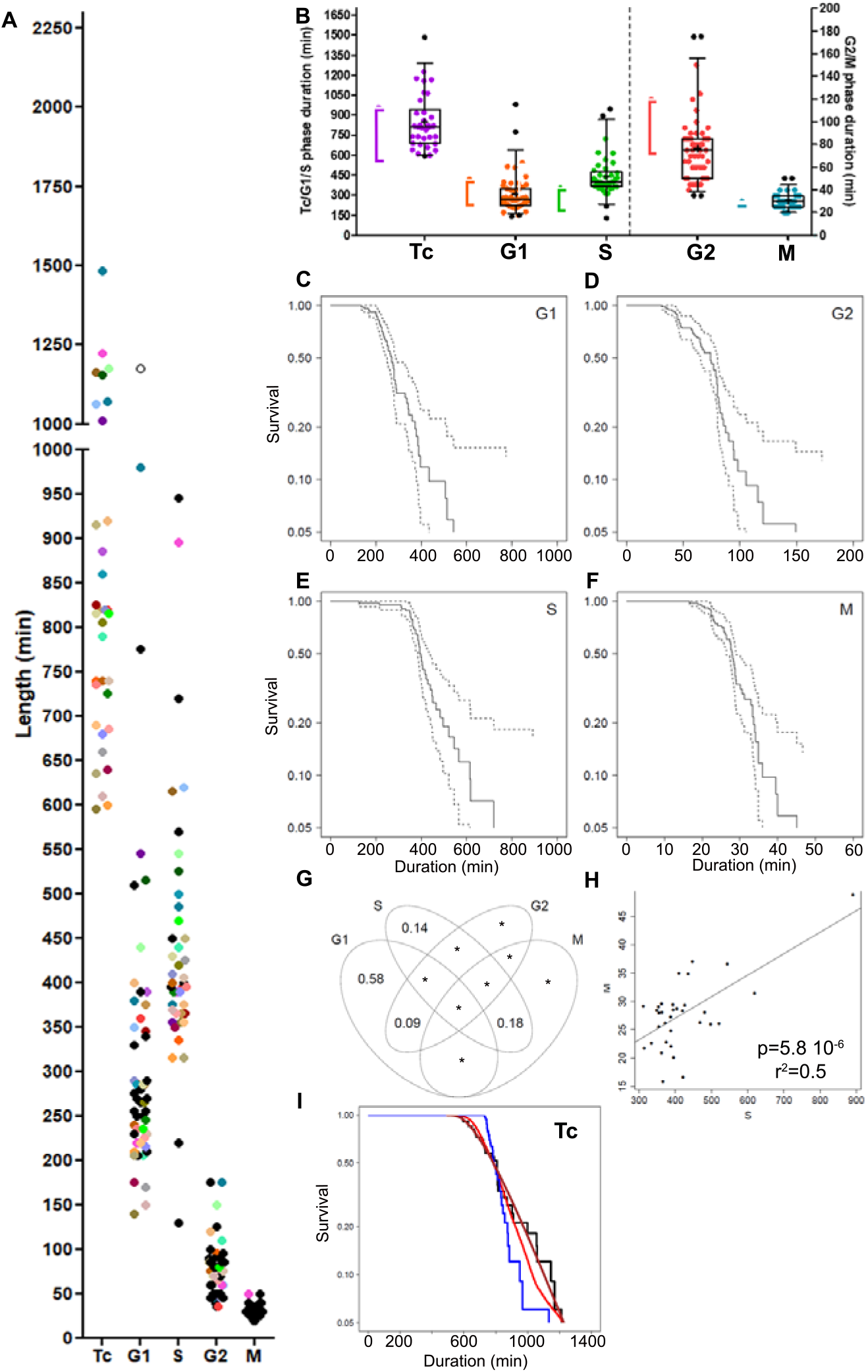
Neural progenitor cell population is highly heterogeneous regarding the cell cycle. **A)** Scatter dot-plot representing distributed lengths of total (Tc) and cell cycle phases for E2.25 embryonic spinal cord cells observed in live imaging. Colored dots in the Tc column can be found in the cell cycle phase columns and correspond to the same tracked nucleus. The empty dot with a G1 length of 1175 min (19h35min) corresponds to a cell that does not start S phase. **B)** Box and whiskers plots (5-95 percentile) illustrating the distribution of total (Tc) and cell cycle phase length from E2.25 embryo cultures and measured in live imaging. Brackets represent values obtained in fixed tissues. The top and the bottom of each box indicate upper and lower quartiles, respectively; the horizontal line represents the median and the cross indicates the mean value. **C-F)** Survival curves to quantify absolute dispersion of G1 phase (**C**), G2 phase (**D**), S phase (**E**), and mitosis (**F**). Black line corresponds to Kaplan-Meier estimates of survival and dashed lines to confidence intervals. **G)** Variation partitioning allow us to examine how much variation of the Tc is attributed to each phase. Stars (*) represent values below 0.05. **H)** Correlation analysis between S and M phases showing a surprising positive coupling. **I**) Tc Survival curve. Black line corresponds to Kaplan-Meier estimate from data. Brown line corresponds to expected survival assuming a random sampling of cell cycle phase duration (see SI-section). Red line corresponds to survival curve obtained in Monte Carlo permutation of phase durations from the data set. Blue curve corresponds to the case with G1 and S/G2/M phase lengths fully anti-correlated (see SI-section).

**Table 1.**

|  |  | Tc (min) | G1 (min) | S (min) | G2 (min) | M (min) |
| --- | --- | --- | --- | --- | --- | --- |
| Control | Mean | 841 | 309 | 434 | 76,6 | 31,1 |
|  | Median | 815 | 268 | 395 | 75 | 30 |
|  | Minimum | 595 | 140 | 130 | 35 | 20 |
|  | Maximum | 1485 | 980 | 945 | 175 | 55 |
|  | Minimal duration (Dm) | 583 | 138 | 128 | 30 | 15 |
| | Exit time ($\tau$) | 247 | 168 | 308 | 44 | 14 |
|  | Std. Deviation | 205 | 148 | 149 | 30 | 7 |
|  | n | 33 | 50 | 41 | 54 | 50 |
| Complete Cycles | Mean | 841 | 307 | 423 | 80 | 30 |
|  | Median | 815 | 265 | 395 | 80 | 30 |
|  | Minimum | 595 | 140 | 315 | 35 | 20 |
|  | Maximum | 1485 | 980 | 895 | 175 | 50 |
|  | Minimal duration (Dm) | 584 | 139 | 311 | 33 | 16 |
| | Exit time ($\tau$) | 247 | 166 | 110 | 45 | 12 |
|  | Std. Deviation | 205 | 156 | 108 | 29 | 5,7 |
|  | Number of values | 33 | 33 | 33 | 33 | 33 |
| PP cells | Mean | 738 | 178,0 | 475,7 | 67,7 | 29,7 |
|  | Median | 610 | 170,0 | 400,0 | 65,0 | 30,0 |
|  | Minimum | 595 | 140,0 | 315,0 | 35,0 | 20,0 |
|  | Maximum | 1225 | 220,0 | 895,0 | 125,0 | 50,0 |
|  | Std. Deviation | 274 | 36 | 202 | 22 | 7,2 |
|  | Number of values | 5 | 5 | 7 | 15 | 15 |
| CDC25B | Mean | 1117 | 494 | 488 | 66,9 | 32,8 |
|  | Median | 1170 | 465 | 493 | 60 | 30 |
|  | Minimum | 595 | 160 | 120 | 10 | 20 |
|  | Maximum | 1585 | 930 | 855 | 170 | 45 |
|  | Minimal duration (Dm) | 586 | 158 | 128 | 30 | 15 |
| | Exit time ($\tau$) | 503 | 333 | 370 | 59 | 14 |
|  | Std. Deviation | 266 | 202 | 148 | 34 | 6 |
|  | n | 23 | 35 | 38 | 61 | 76 |
| CDC25B DCDK | Mean | 809 | 310 | 416 | 79 | 31,5 |
|  | Median | 775 | 265 | 415 | 70 | 30 |
|  | Minimum | 515 | 130 | 105 | 30 | 20 |
|  | Maximum | 1200 | 840 | 930 | 170 | 45 |
|  | Minimal duration (Dm) | 509 | 129 | 103 | 28 | 16 |
| | Exit time ($\tau$) | 291 | 178 | 311 | 49 | 14 |
|  | Std. Deviation | 170 | 145 | 116 | 33 | 5,0 |
|  | n | 45 | 71 | 66 | 86 | 98 |
| E3 | Mean | 865 | 534 | 458 | 70,8 | 32,3 |
|  | Median | 910 | 355 | 480 | 67,5 | 30 |
|  | Minimum | 435 | 180 | 145 | 15 | 25 |
|  | Maximum | 1210 | 965 | 920 | 165 | 45 |
|  | Std. Deviation | 303 | 366 | 203 | 38 | 4,9 |
|  | n | 5 | 10 | 19 | 24 | 24 |

At the single cell level, our analyses showed a high degree of heterogeneity regarding Tc duration, as it ranges from 9h55 min (595 min) to 24h45 min (1485 min) (Table 1; Fig. 3A, B) recalling previous time lapse measurements reporting a total cell cycle length between 9 hours and 28 hours (see FigS2 in (Wilcock, et al., 2007). The use of the FUCCI G1-PCNA biomarker allowed us to determine how this heterogeneity was translated at the level of the different cell cycle phases (Table 1; Fig. 3A). G1 phase durations were comprised between 2h20 (140 min) and 16h20 (980 min). S phase durations spanned from 2h10 (130 min) to 15h45 (945 min), the G2 phase from 35 min to 2h55 (175 min), and Mitosis from 20 min to 55 min. Since data are durations before exiting the phase, we characterized the corresponding distributions using survival analysis (Fig. 3C-F). Each phase can be characterized by a minimal duration (D_min_) and a mean exit time (τ) corresponding to the average duration spent in the phase after that minimal duration (Table 1, SI-Data section 1.2). Mean exit time reflects the slope of the survival function after minimal duration. The gentler the slope, the larger the mean exit time, the more heterogeneous the distribution. At the opposite, a very homogeneous population would display durations close to the minimal duration and a vanishing mean exit time. Hence, the mean exit time is a meaningful readout of the phase duration heterogeneity. The shapes of the survival curves and their mean exit times indicate that all the phases are heterogeneous (Table 1, Fig. 3C, E, SI-Data section 1.3).

To identify the quantitative relationships between phases, we performed further analyses using the subset of 33 cells for which a complete cell cycle was monitored (Fig. 3G, Table 1, SI-Data section 2). In this sample, NPCs cells spent 37% of the cell cycle in G1, 52% in S phase, 9% in G2 and 4% in mitosis, in accordance with previous data obtained on fixed samples (Saade et al., 2013). Variation partitioning analysis showed that 58 % of Tc variation is due to G1 heterogeneity alone (Fig. 3G, SI-Data section 2.2). We then tested correlations within each pair of phases, and found no patterns, except for an unexpected significant positive correlation between S and M durations (Fig. 3H). No correlation was identified between G1 length and S/G2/M lengths (P>>0.05, SI-Data section 2.3.1), suggesting that phases durations are independent from each other within the same cell cycle.

To further test the likelihood of the hypothesis of full independence among phases durations, we examined how it would correctly predict the observed distribution of Tc. To build this prediction analytically, the survival curves of exit times for each phase in the subset have been approximated by a simple exponential decay (red line in Suppl Fig. 2, SI-Data section 2.5.1), which would correspond to a model in which cell cycle phase exit is a stochastic exit process driven by a constant probability per unit time to exit after minimal duration. Under such a hypothesis, for each phase, duration can be described by a stochastic exit process D defined as

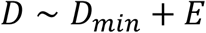

where

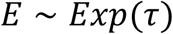

is an exponentially distributed variable with mean time τ.

Under this hypothesis, the complete cell cycle duration would then obey a stochastic process Dc simply defined as:

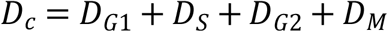

that is, the total duration would just result from the sum of four independent exit processes as defined above, each with its corresponding parameters for minimal duration and mean time before phase exit (see SI-Data section 2.4.1 for the analytical expression for Dc distribution under this hypothesis). We found that the predicted survival function for Dc under this hypothesis (brown curve in Fig. 3I) is actually well compatible with the observed one (black line in Fig. 3I) suggesting that the phases are independent. This is confirmed using Monte Carlo permutation, a random sampling technique (red line in Fig. 3I, SI-Data section 2.4.2). To challenge this finding, we illustrate the opposite hypothesis where Tc length would result from the dataset reordered so that the lengths of G1 and S/G2/M phases are fully anti-correlated (blue line in Fig. 3I, SI-Data section 2.4.3). In this case, the heterogeneity of Tc is greatly reduced compared to the experimental one.

Together these data show that the spinal NPCs population is highly heterogeneous regarding the distribution of cycling time. Duration of each phase seems largely stochastic and independent within one cell cycle, suggesting the absence of coupling between phases. The phase mostly responsible for Tc variation is the G1 phase.

### Maturation correlates with individual NPC G1 phase lengthening

Long-term time-lapse imaging allow one to perform lineage tracing at the single cell level and determine whether a link exists between cell cycle kinetics of mother cells and the fate of daughter cells after mitosis. A proliferating progenitor (P) is characterized when after mitosis it performs a complete G1 phase and enters into S phase. A cell which stop cycling (G1>1000 min; open dot in Fig. 4A) is referred as PLG1. We were able to correctly identify the fate of 15 pairs of daughter cells, 14 being P-P and 1 P-PL_G1_. We then attempted to backtrack the cell cycle features for the corresponding 15 mother cells and measured the duration of 15 mitoses and G2 phases, 7 S phases and 6 G1 phases (colored dots in Fig 4A). Identification of the 14 PP divisions allowed us to compare their cell cycle characteristics to the rest of the population. Regarding the S, G2 and M phases, there is no difference between the distribution of the PP-performing cells and the whole population (p-values of 0.973, 0.141 and 0.215 for S, G2 and mitosis, respectively). However, when considering the G1 phase and Tc lengths, they were significantly decreased by 42.3% and 12.3%, respectively, compared to the phase length measured in the total population (p = 0.001 and 0.041, for the G1 and Tc respectively, Table 1). This is clearly illustrated by the localization of PP-generating mother cells within the lower quartile for the Tc and G1 phase (red dots in Fig. 4A). Extending this analysis to all the PP divisions observed in our various experimental conditions (Suppl. Fig. 3A) confirmed that PP divisions are most often associated with short G1 phase (14 out of 20). Noteworthy, PP division can still be observed with long G1 phase (Suppl. Fig. 3A) suggesting that a long G1 phase does not preclude proliferative division.

**Figure 4.**
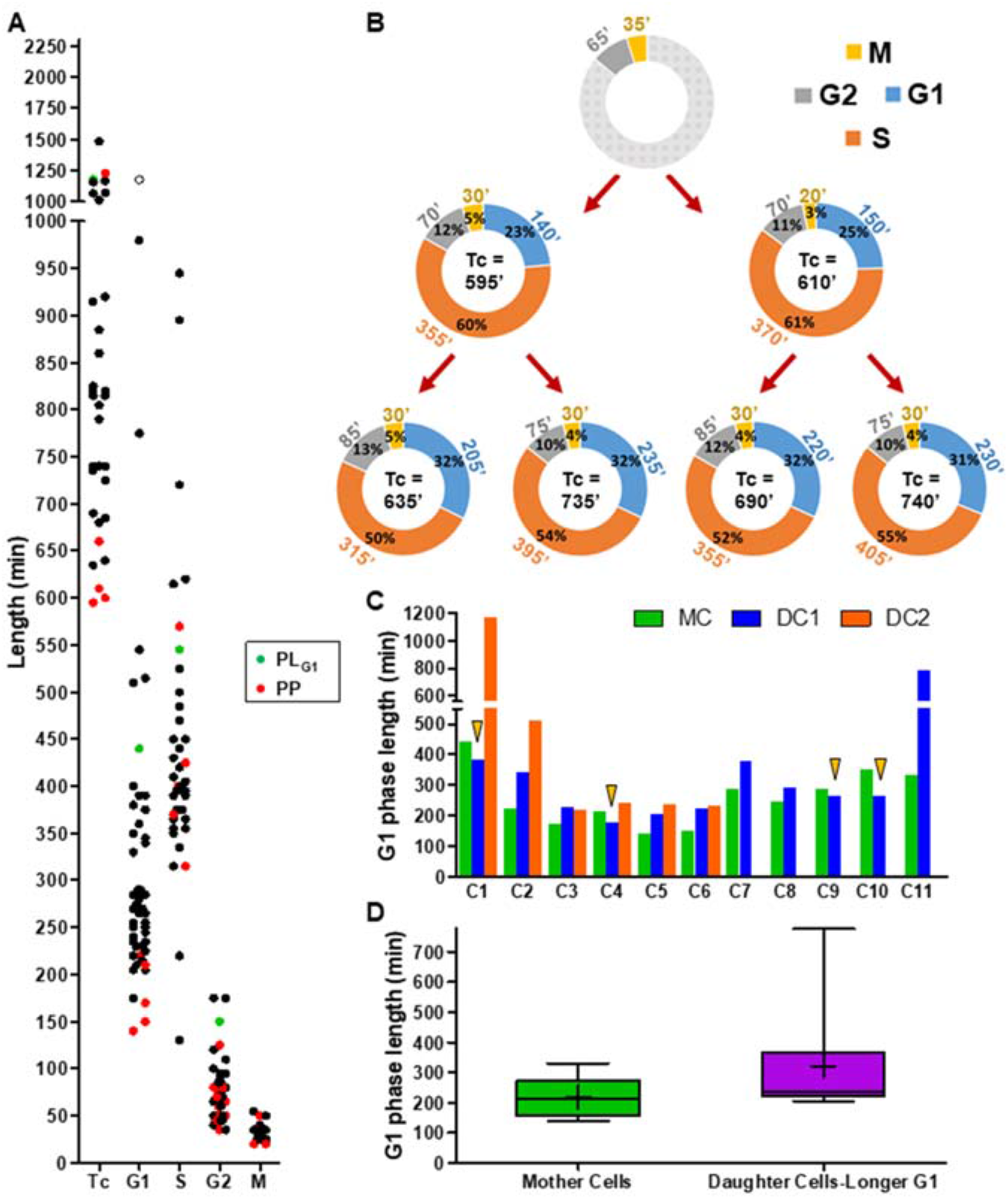
G1 phase lengthens in daughter cells. **A)** Scatter dot-plot representing distributed lengths of cell cycle phases for proliferating NPCs measured in live imaging E2.25 cultures linked to cell fate. Red dots represent NPCs performing PP divisions in the following cell cycle, green dots represent a NPC that gives rise to a progenitor and a cell with a long G1 (asymmetric division), and black dots corresponds to NPCs with undetermined offspring. The empty dot corresponds to a cell with a G1 length of 1175 min (19h35min) that does not start S phase. **B)** Schematic representation of Nuclei tracking over two generations of proliferating NPCs. Cell cycle total (Tc) and phase lengths (G1 in blue, S in orange, G2 in grey and M in yellow) are calculated from time-lapse images. Proportion of each phase in the whole cell cycle according to Tc is represented inside the circles. Light grey represents undetermined values. **C)** Bar plot illustrating G1 phase length of mother (green) and daughter (blue and orange) NPCs. Arrowheads point to daughter cells with a shorter G1 phase as compared to their mother. **D)** Box and whiskers plot representing distributed lengths of G1 phase for mother NPCs and daughter NCPs displaying a longer G1 phase than their mother cell. The top and the bottom of each box indicate upper and lower quartiles, respectively; the horizontal line represents the median and the cross indicates the mean value. n = 8 mother NPCs and 12 daughter cells.

Further lineage analyses revealed that mother and daughter cells displayed differential behavior in terms of cell cycle kinetics. One example of lineage is given in Fig. 4B). For the G1 phase, 13 out of 17 daughter cells (76.5%) presented a longer G1 phase than their mother without difference in the S, G2, and Tc cell cycle phases (Fig.4C). The mean difference between these “longer G1 daughter cells” and their mothers is of 1h42 min (Fig. 4D). The 4 other cells (arrowheads in Fig. 4C) display a shorter G1 phase with a reduction of 50 min (Suppl. Fig. 3B). Our results suggest that G1 phase lengthening occurs from generation to generation.

If our statement is true, at later developmental stages when the population of neurogenic NPCs increases at the expense of progenitors performing PP divisions, we should observe a modification in the distribution of the G1 phase length. We therefore electroporated E2 embryos, waited for 24 hours before processing them for live imaging (Fig. 1E) and performed cell cycle live imaging in E3 (HH20) embryonic neural tube explants. As observed in Fig. 5A, out of the 30 G1 nuclei analyzed, 20 display a G1 phase longer than 1000 min after mitosis (open circle in Fig. 5A) as expected from a population of NPCs containing numerous progenitors committed to neuronal differentiation. The other 10 cells performed a complete G1 phase and start an S phase. The total cell cycle length was measured on 5 out of these 10 cells (colored dots in Fig. 5A, Table 1). The Tc, S, G2 and M phases showed similar mean values with those measured at E2 while the average duration of G1 phase was increased (8h54 at E3 vs 5h09 at E2; p = 0.316). The frequency distribution comparison between E2 and E3, revealed for G1 phase, the appearance of cells with G1 around 900 min creating a second peak on the left of the figure (Fig. 5B). S phase data show an asymmetrical skewed distribution with a difference in the peak position and data spreading revealing a right shift in the distribution indicating a lengthening and a left shift in G2 phase representing a shortening (Fig. 5C-5D). Unfortunately, out of 10 cycling G1 nuclei, we were only able to backtrack the cell cycle of one mother cell whose two daughters performed long G1s (blue dot in Suppl. Fig. 3A). Altogether these data suggest that NPCs performing PP divisions will mainly execute a short G1 phase. G1 lengthening from generation to generation is correlated with NPCs maturation and possibly neuronal fate commitment.

**Figure 5.**
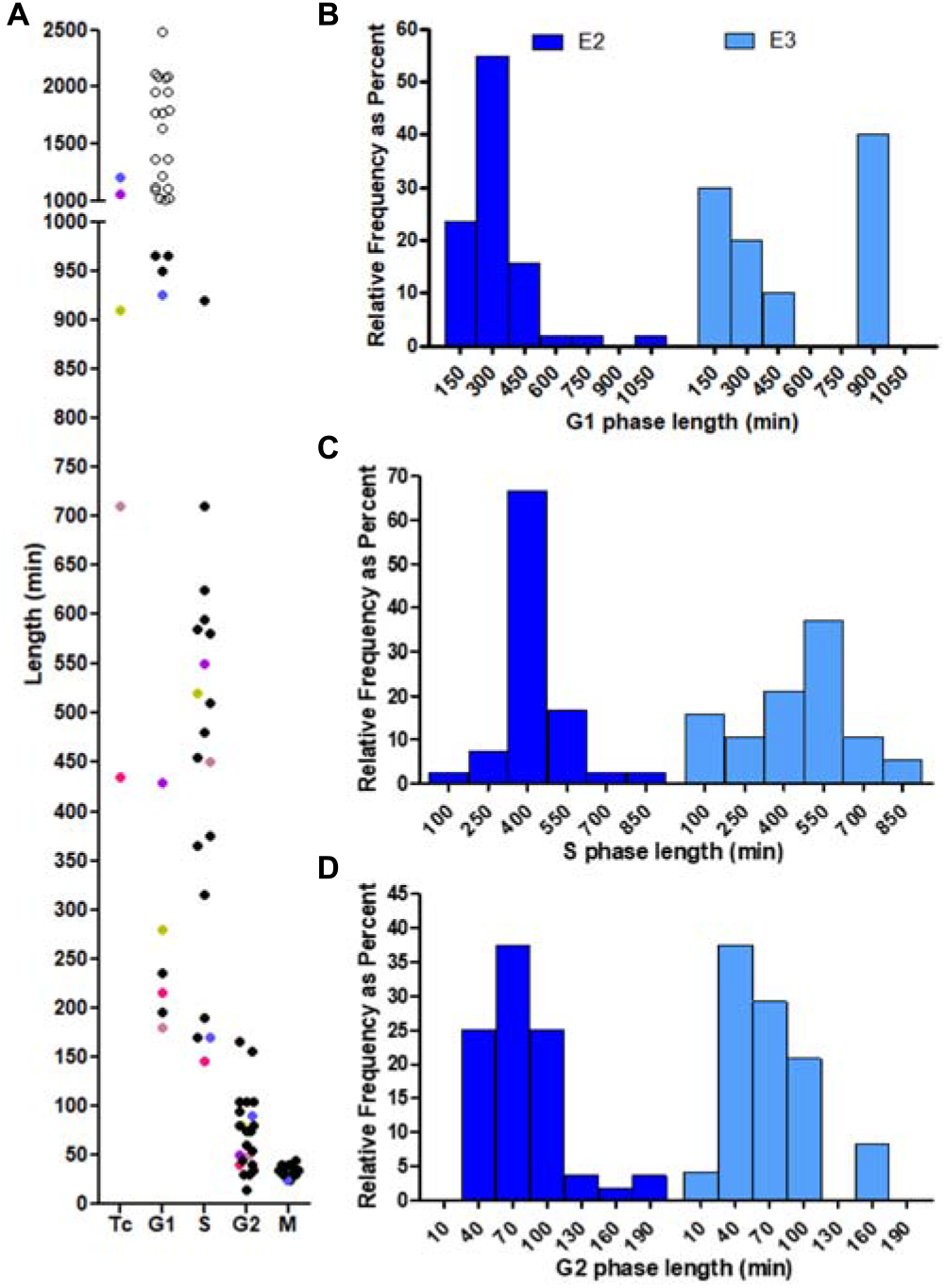
Neural progenitor cell population kinetics changes in older embryos. **A)** Scatter dot-plot representing distributed lengths of total (Tc) and cell cycle phases for embryonic spinal cord nuclei measured from E3 spinal cord cultures live imaging. Colored dots in the Tc column can be found in cell cycle phases columns and correspond to the same tracked nucleus. Empty dots represent cells with G1 phase length superior to 1000 min and correspond to those cells that do not start S phase. **B-C-D)** Frequency distribution of G1 (**B**), S (**C**) and G2 (**D**) phase lengths comparing E2.25 spinal cord cultures (dark blue) and E3 spinal cord cultures (light blue).

### CDC25B activity induces G1 phase lengthening in a CDK dependent manner

As mentioned above, single cell analysis reveals a high degree of heterogeneity in G1 phase length at the population level. Lineage experiments indicate that daughter cells display most often a longer G1 phase than their mother suggesting that G1 phase lengthens hereditarily. One possibility is that a mechanism intrinsic to the cell cycle induces a lengthening of the G1 phase in individual NPCs. Recently, we showed that CDC25B is involved in NPCs maturation (Bonnet, et al., 2018; Peco, et al., 2012), we then decided to test the effects of the phosphatase on cell cycle kinetics using our time lapse strategy. We performed gain of function using a vector that reproduces the iterated cell cycle regulated expression of CDC25B (Bonnet, et al., 2018). As observed in control condition, cell cycle kinetics is highly heterogeneous in CDC25B gain-of-function (Fig. 6A and Table 1). As expected from its role in G2/M transition (Bonnet, et al., 2018; Peco, et al., 2012), CDC25B gain of function induced a significant 12.7% decrease of the mean G2 phase length (p = 0.029, Fig. 6C and Table 1). This effect on G2 is associated with a mean Tc length increase of 32.8% (p=0.004 when compared to control; Table 1) resulting from a slight increase in the length of the mean S and M phase lengths (12.4% and 5.5% respectively) and from a drastic 59.9% increase in the mean G1 phase length (p<0.0001 when compared to control; Fig.6C, Table 1). Survival curves and the corresponding frequency distribution histograms representation were then compared to analyze the dispersion of the data in various conditions (Fig. 6D-H). The 4h16 min (256 min) increase observed in mean Tc length induced by CDC25B is associated with a rise of the exit time (from 247 to 503 min) without modification of the minimal duration (583 min vs 586 min, Table 1). This clearly define that CDC25B increases the dispersion of the dataset. The pace at which CDC25B expressing cells exit the cell cycle is about three times slower (hazard ratio of 0.35) than that of control cells (red curve compared to black curve in Fig. 6D, SI-Data section 3.1). When comparing phase by phase, CDC25B induces a significant increase in both G1 and S phase distribution compared to control (compare red and black curve in Fig. 6E-G). Minimal durations do not appear altered, but rather exit times are affected, with a mean lengthening of 2h45 (165 min) and 1h02 (62 min) and 0.41 and 0.65 hazard ratios for G1 and S phases, respectively (SI-Data section 3.2 and 3.3). No significant effect was found on the G2 or M phase survival distributions (Fig. 6F-H, Table 1, SI-Data section 3.4 and 3.5). We note that mean G2 length shortening appears to be associated with a shorter minimal duration, with no effect upon its heterogeneity (shape of the curve in Fig. 6 F, Table1). These data suggest that CDC25B affects primarily G1 phase heterogeneity.

**Figure 6.**
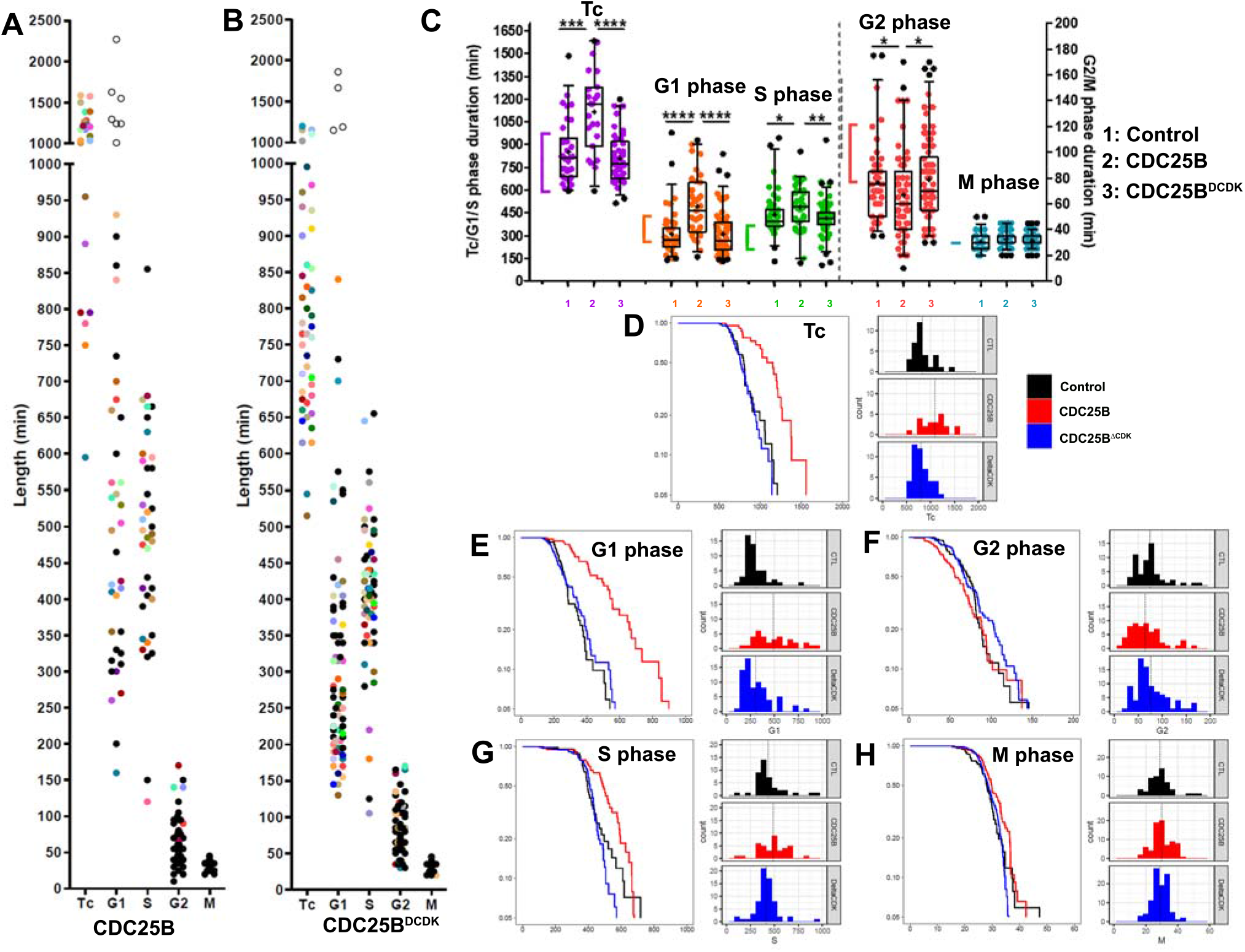
CDC25B gain-of-function increases G1 phase length heterogeneity. **A-B)** Scatter dot-plot representing distributed lengths of total (Tc) and cell cycle phases measured in live imaging E2.25 cultures for cells electroporated with CDC25B (**A)** or CDC25B^ΔCDK^ (**B).** Colored dots in the Tc column can be found in the cell cycle phase columns and correspond to the same tracked nucleus. Empty dots represent cells with a G1 length longer than 1000 minutes (16h40min) that do not start S-phase. **C)** Box and whiskers plots (5-95 percentile) illustrating the comparison of total (Tc) and cell cycle phase lengths for Control, CDC25B, CDC25B^ΔCDK^ gain-of-function. Brackets represent values in fixed tissues. The top and the bottom of each box indicate upper and lower quartiles, respectively; the horizontal line represents the median and the cross indicates the mean value. (**D-H**) Survival curves and histogram representation comparing Control (black), CDC25B (red), and CDC25B^ΔCDK^ (blue) conditions for the Tc (**D**), G1 **(E**), G2 **(F**), S **(G**), and M (**H**), phases. The vertical dotted line in histograms reports the average of the distribution.

Variation partitioning using complete cell cycle revealed that about 52 % of Tc variation are explained by G1 heterogeneity alone and 21% by S phase alone in CDC25B condition (Suppl. Fig. 4A). In this condition, correlation analysis between phase lengths showed that phases are uncoupled (SI-Data section 2.3.2). Monte Carlo permutations of the data set (10^4^ samples) suggest that the distribution of Tc is compatible with independent cell cycle phase durations (compare red and black curve in Suppl. Fig. 4B and SI-Data section 2.6.2). Because CDC25B effectors control directly G2 phase length, we tested for a correlation between the mother cells G2 phase length and the daughter cells G1 phase length. We could not detect a heritable correlation neither in control nor in the gain of function condition (Suppl. Fig. 4C, D).

We then wanted to know whether the effects of CDC25B on the G1 phase was dependent on the interaction with CDK. We electroporated a CDC25B^ΔCDK^ mutant form of CDC25B unable to interact with CDK but still displaying neurogenic activity albeit less important than the wild type protein (Bonnet, et al., 2018). In line with the neurogenic effect of this mutant protein, we observed 4 cells with a G1>1000 minutes in this condition (open circles in Fig. 6B). In our experiments, CDC25B^ΔCDK^ neurogenic mutant was not able to significantly change cell cycle phases length and dispersion compared to control conditions (Fig. 6B, C; Table 1, SI-Data section 3). Thus, the G1 phase modification induced by CDC25B is dependent on its interaction with CDK.

All these results show that in a context where each cell cycle phase duration is stochastic and independent, CDC25B enhances heterogeneity in G1 phase duration leading to a lengthening of G1 and Tc mean values at the population level, which is reminiscent of what is observed in maturing neural tissue.

## Discussion

In this paper we detailed a live imaging strategy allowing us to follow the behavior of single NPCs over 48 hours in their endogenous environment. We show that the cell cycle duration and, more interestingly, the length of each phase is very heterogeneous and with no apparent links between phase lengths within one cell cycle or between mother cells G2 phase and daughter cells G1 phase. Though, we find that G1 phase length increases with cell generations and is the one which contributes mainly to the total cell cycle lengthening. We show that expression of the G2/M regulator CDC25B enhances G1 length heterogeneity in NPCs participating to neural tissue maturation.

### Neuroepithelial single cell imaging of the cell cycle in its endogenous environment

Measuring the four phases of the cell cycle in real time with a single cell resolution within a tissue remains challenging. Here, we performed *ex ovo* live imaging using chick embryo slice cultures (Das, et al., 2012; Pituello, et al., 1995). In our hands, three parameters are then critical: cell cycle biosensor, culture conditions and imaging methods. Dynamic analysis of cell cycle progression by real-time imaging has become feasible with the generation of the FUCCI reporter system (Sakaue-Sawano, et al., 2008) recently improved for the detection of the four phases (FUCCI4, (Bajar, et al., 2016). We did not favor the FUCCI4 strategy because the increased light illumination required for four laser illumination can be detrimental to cell survival. Our biosensor assay combines the mKO-zCdt1 and the NLS-eGFP-PCNA discriminates the four phases of the cell cycle with only two different wavelengths illuminations per time point. Another critical point is the culture conditions. To ensure cell survival and obtain cell cycle parameters comparable to those measured on fixed tissue, it is important to add 5% FCS and to allow tissue recovery for 12 hours. In such conditions NPCs perform proliferative and neurogenic divisions. Finally, time lapse imaging systems are also critical for cell survival and normal cell behavior. We observed that not only limiting illumination by exciting a focal plane (wide field vs confocal microscope) but also acquisition velocity (confocal vs spinning disk equipped confocal microscope), were crucial to decreasing cell death by phototoxicity (as also described by (Icha, et al., 2017). Similarly, laser beam pulses frequencies play a role in cell cycle kinetics and survival, this is why we opted for a spinning disk system with laser attenuation (Boudreau, et al., 2016) to reduce the stress to the cells. We nevertheless observed a slightly longer S phase as already described in another study (Leung, et al., 2011).

### The lengthening of the cell cycle results from enhanced heterogeneity

We show that the total cell cycle length exhibits a high degree of heterogeneity, ranging from 9h55min to 24h45 min without apparent patterns or coupling between the phase length except a surprising link between the S and M phase. Such a heterogeneity of the total cell cycle length has already been observed using time lapse imaging in neural stem/progenitor cells including spinal NPCs (Wilcock, et al., 2007), mice neural stem cells in culture (Roccio, et al., 2013), or human nervous system primary tissues and organoids (Subramanian, et al., 2017). This cell cycle heterogeneity is also observed at different levels for each phase of the cell cycle, the phase that contributes the most to the total cell cycle variability being the G1 phase. Usually, analyses of cell cycle parameters were performed on fixed tissues, considering that neural stem cells are a homogenous population of asynchronous proliferating cells. Using this approximation, these analyses showed that as developmental time progresses, the proliferation rate of neural progenitors decreases and their cell cycle lengthens (Molina and Pituello, 2017; Kicheva and Briscoe, 2015; Kicheva, et al., 2014). This evolution of the cell cycle length most often associated with differentiation was also observed in various stem cell types (Julian, et al., 2016; Dalton, 2015). If the population of NPCs was homogeneous, an increase of the mean cell cycle duration with time would correspond to an increase of the cell cycle duration in each individual NPCs. Instead, we observed that the population of NPCs is heterogeneous, and in this case the increase in the mean cell cycle duration occurring with time correlates with an increase in heterogeneity. To test possible relationships between the four phase lengths, we developed a mathematical model in which Tc length results of the sum of cell cycle phase duration stochastic and independent of each other. The experimental and theoretical Tc survival curves display a very similar pattern, suggesting that indeed cell cycle phase durations are stochastic and independent. One hypothesis is that permitting such stochastic variations at the single cell level is cost efficient for the population, which does not need to strictly control the cell cycle kinetics of each single cell and thereby is probably more robust. These results are reminiscent of experimental data and modeling obtained using NIH3T3 cell cultures that showed that lengthening one phase does not imply lengthening other phases, suggesting that durations of cell cycle phases can be regarded as independent (Mura, et al., 2019). This is also observed for three independent human cell lines, where each cell cycle phase has been shown to be independent of the others and that cell cycle correspond to a series of uncoupled memoryless phases (Chao, et al., 2019).

We propose that i) the total cell cycle length in NPCs corresponds to addition of independent and stochastic phase lengths, ii) the increase of cell cycle length observed during development, is due to an increased heterogeneity of cell cycle length in the NPCs population. This suggests that there is no regulatory system that controls connections between each phase length and that a mechanism increasing cell cycle heterogeneity might be responsible for the modifications observed during stem cell maturation. In the mouse epidermal stem cells, a pronounced heterogeneity of cell cycle length has been recently described using time-lapse (Xie and Skotheim, 2020). Their analysis of the 3D volume growth and cell-cycle progression of single epidermal stem cells showed that cell cycle length is controlled by cell growth through a sizer operating in the G1 phase of the cell cycle (Xie and Skotheim, 2020). In this case, cell growth heterogeneity would control cell cycle heterogeneity. We can imagine that a similar mechanism takes place in the neuroepithelium. One way to control cell size is to affect cell metabolism, and several links have been described between modification of metabolic pathways and modulation of neurogenic activities during adult neurogenesis (Knobloch and Jessberger, 2017) or embryonic neurogenesis (Fawal, et al., 2018).

### A novel mechanism to control the G1 and cell cycle lengthening

Time lapse analysis showed that even if all cell cycle phase lengths are heterogeneous the S and the G1 phase contribute the most to the total cell cycle heterogeneity, and that the G1 phase is the most affected one upon development. G1 phase heterogeneity has already been observed in neuroepithelial cells in culture (Roccio, et al., 2013), in human embryonic stem cells in culture (Jang, et al., 2019), and in mouse epidermal stem cell in vivo (Xie and Skotheim, 2020). Here we identified a new trigger of the G1 phase lengthening, the CDC25B phosphatase. CDC25B proteins have long been known to regulate G2 phase to mitosis transition by activating CDK1/cyclin B complexes (Kumagai and Dunphy, 1991). We have shown that the expression of the phosphatase in NPCs correlates spatially and temporally with neurogenesis (Bonnet, et al., 2018; Agius, et al., 2015; Peco, et al., 2012). In agreement with this expression pattern, using both gain and loss of function, we showed that CDC25B is a positive regulator of neurogenesis (Bonnet, et al., 2018; Peco, et al., 2012) and contributes to neural tissue aging (Azaïs, et al., 2019; Bonnet, et al., 2018).

Part of CDC25B neurogenic activity is independent of its effect on the cell cycle, however, its full neurogenic potential requires a cell cycle dependent action (Bonnet, et al., 2018). Here we show that CDC25B induces a lengthening of the mean G1 phase duration in a cell cycle dependent manner. It acts by increasing G1 heterogeneity, as illustrated by the fact that exit time is almost doubled and exit rate is more than twice slower than control. The full neurogenic activity of CDC25B will then be the sum of its ability to induce neurogenic division independently of CDK interaction and its action as a maturating factor reducing the proliferative capacities of the NPCs.

Electroporation in chick neural tube leads to mosaic expression. We therefore considered whether part of the G1 heterogeneity observed results from the method we used. The primary activity of CDC25B is to control G2/M transition and thereby affects G2 phase length. In our experiment, the effect observed on G1 heterogeneity is associated with a reduction of the G2 minimal duration in the absence of an effect on the G2 phase heterogeneity (compare the slope of the survival curves, see Fig. 6E and 6F). This suggests that the mosaic expression induced by electroporation, does not increase heterogeneity in a non-specific way.

Lengthening of the G1 phase has long been associated with neurogenesis (Kuzmicz-Kowalska and Kicheva, 2020; Molina and Pituello, 2017; Takahashi, et al., 1995). Multiple molecular mechanisms link G1 phase length and choice between proliferation and differentiation in multiple stem cell types, including various pluripotent stem cells (Julian, et al., 2016; Dalton, 2015) and neural stem cells (Liu, et al., 2019). In the mouse cortex, overexpression of cyclin D1, cyclin E1, or CDK4, increases self-renewal and inhibits neurogenic differentiation, establishing a link between G1 length and neurogenesis (Lange, et al., 2009; Pilaz, et al., 2009). In the spinal cord, forced expression of G1-phase regulators (D-type cyclins) inhibited the neural marker Tuj1 expression 24 hours after electroporation (Lacomme, et al., 2012). Conversely, NSCs of Cdk2/Cdk4 double-knockout mice displayed an increased propensity for neuronal differentiation, leading to enhanced neurogenic divisions in the brains of the mutant embryos (Lim and Kaldis, 2012). Thus, by lengthening the G1 phase, CDC25B might create conditions for differentiation.

What is the link between G1 phase length and cell fate? In hESC it has been shown that differentiation capacity varies during progression of the G1 phase (Pauklin and Vallier, 2013). In this case, G1 lengthening has been associated with modifications of TGFβ signaling cascades and ectoderm vs endo mesoderm cell fate choice. They showed that Smad2,3 activates mesoderm and endoderm commitment in early G1, and that ectoderm commitment occurs in late G1, coinciding with nuclear exclusion of SMAD2,3 in a Cyclin D dependent manner. Alternatively, CDKs may regulate by phosphorylation the activity of transcription factors involved in proliferation or differentiation. For example, CDK-dependent phosphorylation of Sox2 at Serine 39 inhibits neurogenesis, and CDK decrease leads to proteolytically cleaved Sox2 species that promotes neurogenesis (Lim, et al., 2017). In addition, cyclin A- and B-dependent kinases phosphorylate the proneural basic helix-loop-helix transcription factor Neurogenin 2 (Ngn2) and inhibit its ability to bind neurogenic genes and to promote neurogenesis (Ali, et al., 2011). Interestingly, Ngn2 can repress expression of cyclins D1 and E2, suggesting the presence of a positive-feedback loop (Lacomme, et al., 2012). In CDC25B gain of function experiments, the total cell cycle duration was increased at least partly as a consequence of G1 phase lengthening. In our recent theoretical studies (Azaïs, et al., 2019), we proposed that CDC25B expression in neural progenitors progressively restricts proliferative capacities of the cell. We propose that CDC25B reiteration at each cell cycle will indirectly increase G1 phase heterogeneity, lengthening cell cycle duration associated with differentiation and participating to tissue maturation. Such a mechanism is likely to be applicable to other developing organs or tissues as well as to stem cells including human stem cells

## Acknowledgement

We want to thank Bernard Ducommun, Alice Davy and Xavier Morin for critical review of the manuscript. Caroline Monod for improving the English. We thank the CBI Toulouse Regional Imaging platform (TRI) for technical support. We acknowledge the Developmental Studies Hybridoma Bank, created by the NICHD of the NIH and maintained at The University of Iowa, Department of Biology, Iowa City, IA 52242, for supplying monoclonal antibodies. Work in FP’s laboratory is supported by the Centre National de la Recherche Scientifique, Université P. Sabatier, Ministère de L’Enseignement Supérieur et de la Recherche (MESR) et Agence Nationale de la Recherche (ANR-19-CE16-0006-01). Angie Molina was a recipient of IDEX UNITI and Fondation ARC. Fréderic Bonnet was recipient of MESR studentships. The funding entities had no role in study design, data collection and analysis, decision to publish, or preparation of the manuscript.

## Author Contributions

A.M., F.B., F.P., J.G. and E.A. conceived experiments. A.M., F.B., and E.A. performed experiments. A.M., F.B., F.P. J.G. and E.A wrote the manuscript. F.P., J.G., V.L. and E.A secured funding. S.B.V and V.L. provided expertise and feedback

---

**Supplement Figure 1. Characterization of *ex vivo* culture of chick embryonic neural tube explants. A, B)** Cross-sections of E2.5 and E3.5 (stage HH18, HH23) chick embryo spinal cord (in ovo), and explants dissected at E1.5 and cultivated for 24 hours and 48h hours (ex ovo). Sections processed for anti-caspase3 (green) and anti-MNR2 (red) immunostaining (**A**) or anti-Olig2 (red) and anti-lslet1/2 (green) immunostaining (**B**). Inset in B, E1.5 (HH13) embryo section showing olig2 progenitors. **C)** Scatter dot plot representing S phase duration (Ts) and total cell cycle duration (Tc) calculated using Dual Pulse Labeling using EdU and BrdU incorporation paradigm in embryo and in culture, revealing a transient lengthening of the cell cycle in progenitors at 12 hours after dissection that is recovered at 24 hours (not shnow). **D)** Curves representing kinetics of the number expressing Olig2 and lslet1/2 per section in the spinal cord (in ovo) or in explants (ex vivo) starting at E1.5 (t = Oh), 24 hours or 48 hours later. The increase in the population of progenitors and neurons in our culture conditions indicates that progenitors are performing both proliferative and neurogenic divisions. Data from three different experiments with at least four embryos for the control, and six sections from three embryos for the cultures in each condition. Scale bars represent 100 μm.

**Supplement Figure 2**. Survival curves for exit times for the four phases, from the data subset of complete cell cycle tracked cells. Exit times are obtained by subtracting minimal time from observed times. Black curves are Kaplan-Meier estimates of survival, with confidence interval. For this subset, survival curves appear to be about compatible with simple exponential decay (red curves), corresponding to a simple memoryless process.

**Supplement Figure 3. Modes of division and G1 phase length. A**) Box and whiskers plot representing distributed lengths of G1 phase in Control, CDC25B, CDC25B^ΔCDK^ and E3 experiments associated with Scatter dot-plot representing distributed G1 lengths of NPCs performing PP divisions in the following cell cycle (red dots), NPCs giving rise to a progenitor and a cell with a long G1 (asymmetric division, green dots) and NPCs generating 2 daughter cells with long G1s (blue dots). **B**) Box and whiskers plot representing distributed lengths of G1 phase for mother NPCs and daughter NCPs displaying a shorter G1 phase than their mother cell. The top and the bottom of each box indicate upper and lower quartiles, respectively; the horizontal line represents the median and the cross indicates the mean value. n = 4 mother NPCs and 4 daughter cells.

**Supplement Figure 4. CDC25B effects on cell cycle heterogeneity A)** Variation partitioning (Venn Diagrams) in CDC25B gain of function showing how much variation is attributed to each phase. Stars (*) represent values below 0.05. **B** Survival curve of Tc phase length data in the CDC25B condition. Black line corresponds to Kaplan-Meier estimate from data. Red line corresponds to the survival curve obtained in Monte Carlo permutation of phase durations from the data set and the blue curve corresponds to the case with G1 and S/G2/M phases lengths fully anti-correlated (see SI-sect). **C, D)** Correlation analysis between G2 phase lengths of the mother cell and G1 phase lengths of the daughter cell in control (**C**) and CDC25B (**D**).

**Video 1. Expression of the cell cycle biosensor within the nucleus of NPCs allows the identification of the four cell cycle phases.** Left panel-Live imaging movie. The four cell cycle phases are discriminated by the differential expression of the sensor, as well as the movements of nuclei inside the neural tube (Interkinetic Nuclear Movement, INM). Right panel - Segmentation of some analyzed nuclei. Time interval between frames = 5 min. Scale bar = 10 μm.

