## Supplementary material for "G1 Phase Lengthening During Neural Tissue Development Involves CDC25B Induced G1 Heterogeneity": supp figure

### Figure sup 1

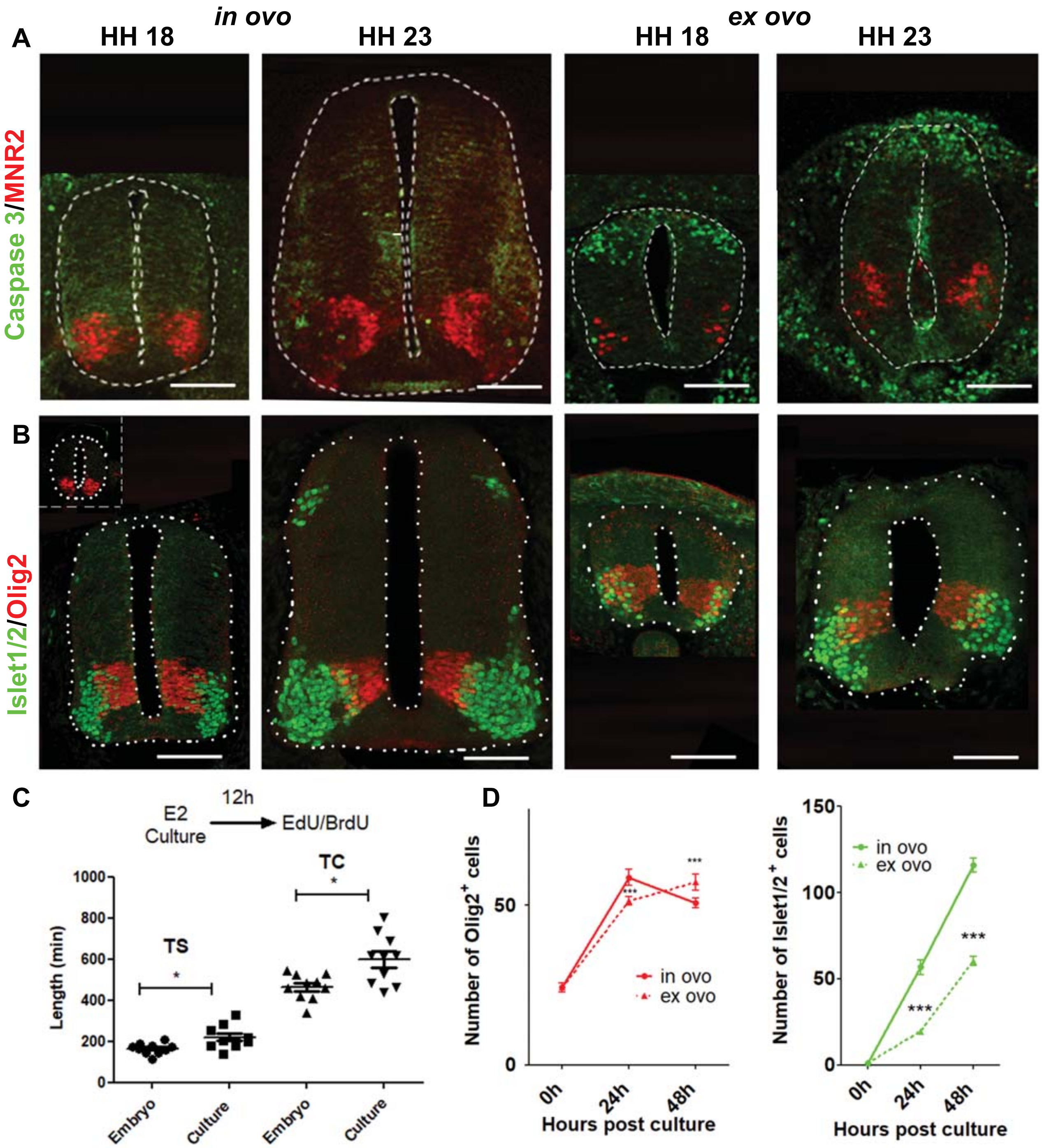

A
in ovo
ex ovo
HH 18
HH 23
HH 18
HH 23
Caspase 3/MNR2
B
Islet1/2/Olig2
C
E2
Culture
12h
EdU/BrdU
TS
TC
Length (min)
1000
800
600
400
200
0
Embryo
Culture
Embryo
Culture
D
Number of Olig2+ cells
50
0
0h
24h
48h
Hours post culture
in ovo
ex ovo
***
Number of Islet1/2+ cells
150
100
50
0
in ovo
ex ovo
***
0h
24h
48h
Hours post culture

### Figure sup 2

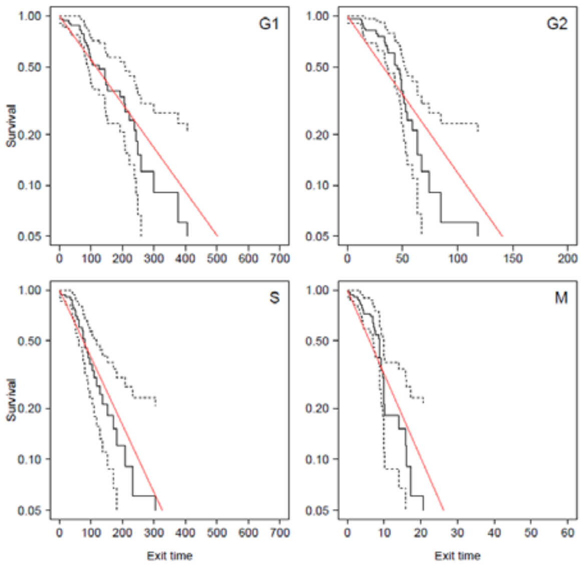
G1
G2
S
M
Survival
Exit time
1.00
0.50
0.20
0.10
0.05
0 100 200 300 400 500 600 700
0 50 100 150 200
0 10 20 30 40 50 60

### Data Analysis For The Article *G1 PHASE LENGTHENING DURING NEURAL TISSUE DEVELOPMENT INVOLVES CDC25B INDUCED G1 HETEROGENEITY*

Molina et al. 2020

### 1 Data Descriptions

We read durations data from 00-AllPhasesData.csv.

For complete cycles, we can compute Tc.

Note : in R outputs all along below, the condition is encoded as factor according to :

| Condition | Code |
|---|---|
| CONTROL | CTL |
| CDC25B | CDC25B |
| CDC25B$^{\Delta CDK}$ | DeltaCDK |

#### 1.1 Data summaries

##### 1.1.1 CONTROL

```
   Condition         G1               S                 G2                M                Tc        
 CTL     :54   Min.   :140.0   Min.   :130.0   Min.   : 35.00   Min.   :20.00   Min.   : 595.0  
 CDC25B  : 0   1st Qu.:227.5   1st Qu.:365.0   1st Qu.: 52.50   1st Qu.:25.00   1st Qu.: 690.0  
 DeltaCDK: 0   Median :270.0   Median :397.5   Median : 75.00   Median :30.00   Median : 815.0  
               Mean   :308.7   Mean   :438.6   Mean   : 76.57   Mean   :31.27   Mean   : 840.9  
               3rd Qu.:347.5   3rd Qu.:465.0   3rd Qu.: 85.00   3rd Qu.:35.00   3rd Qu.: 915.0  
               Max.   :980.0   Max.   :945.0   Max.   :175.00   Max.   :55.00   Max.   :1485.0  
               NA's   :3       NA's   :12                       NA's   :3       NA's   :21      
```

##### 1.1.2 CDC25B

```
   Condition         G1               S                 G2                M                Tc        
 CTL     : 0   Min.   :160.0   Min.   :120.0   Min.   : 10.00   Min.   :20.00   Min.   : 595.0  
 CDC25B  :76   1st Qu.:327.5   1st Qu.:401.2   1st Qu.: 40.00   1st Qu.:30.00   1st Qu.: 906.2  
 DeltaCDK: 0   Median :465.0   Median :492.5   Median : 60.00   Median :30.00   Median :1130.0  
               Mean   :494.4   Mean   :487.5   Mean   : 66.89   Mean   :32.76   Mean   :1099.8  
               3rd Qu.:625.0   3rd Qu.:587.5   3rd Qu.: 80.00   3rd Qu.:35.00   3rd Qu.:1255.0  
               Max.   :930.0   Max.   :855.0   Max.   :170.00   Max.   :45.00   Max.   :1585.0  
               NA's   :41      NA's   :38      NA's   :15                       NA's   :54      
```

##### 1.1.3 CDC25B$^{\Delta CDK}$

```
   Condition         G1               S                 G2                M                Tc        
 CTL     : 0   Min.   :130.0   Min.   :105.0   Min.   : 30.00   Min.   :20.00   Min.   : 515.0  
 CDC25B  : 0   1st Qu.:207.5   1st Qu.:376.2   1st Qu.: 56.25   1st Qu.:30.00   1st Qu.: 680.0  
 DeltaCDK:98   Median :265.0   Median :415.0   Median : 70.00   Median :30.00   Median : 775.0  
               Mean   :309.6   Mean   :416.4   Mean   : 79.13   Mean   :31.48   Mean   : 809.3  
               3rd Qu.:377.5   3rd Qu.:455.0   3rd Qu.: 93.75   3rd Qu.:35.00   3rd Qu.: 910.0  
               Max.   :840.0   Max.   :930.0   Max.   :170.00   Max.   :45.00   Max.   :1200.0  
               NA's   :27      NA's   :32      NA's   :12                       NA's   :53      
```

#### 1.2 Statistics of durations

For each phase: minimal duration, standard deviations of phases duration and mean latency before exiting the phase starting from this minimum (hereafter denoted *Exit times*).

##### 1.2.1 Minimal durations

| Condition | G1 | S | G2 | M | Tc |
|---|---|---|---|---|---|
| CONTROL | 140 | 130 | 35 | 20 | 595 |
| CDC25B | 160 | 120 | 10 | 20 | 595 |
| CDC25B$^{\Delta CDK}$ | 130 | 105 | 30 | 20 | 515 |

##### 1.2.2 Standard deviation of phase durations

| Condition | G1 | S | G2 | M | Tc |
|---|---|---|---|---|---|
| CONTROL | 148 | 149 | 30 | 7 | 205 |
| CDC25B | 202 | 147 | 34 | 6 | 266 |
| CDC25B$^{\Delta CDK}$ | 144 | 116 | 33 | 5 | 170 |

##### 1.2.3 Mean exit times

| Condition | G1 | S | G2 | M | Tc |
|---|---|---|---|---|---|
| CONTROL | 169 | 309 | 42 | 11 | 246 |
| CDC25B | 334 | 368 | 57 | 13 | 505 |
| CDC25B$^{\Delta CDK}$ | 180 | 311 | 49 | 11 | 294 |

##### 1.3 Survival curves for each condition.

We add noise to avoid ties. Actually, snaps were made every 5 min, so the event has occured within the last 5 minutes. Hence we distribube its date randomly within those last 5 minutes.

###### 1.3.1 CONTROL

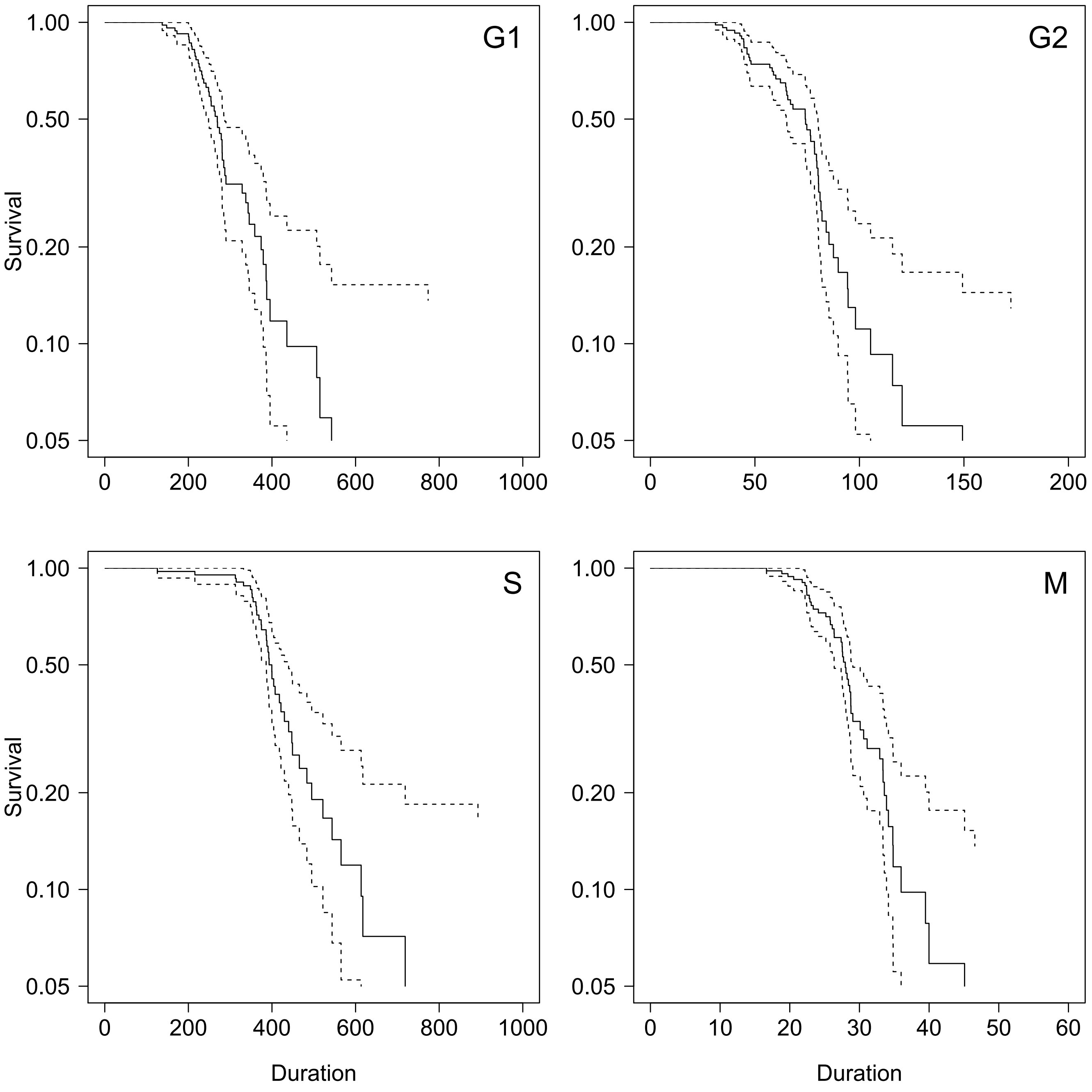

Figure 1: Phases Survival curves

###### 1.3.2 CDC25B

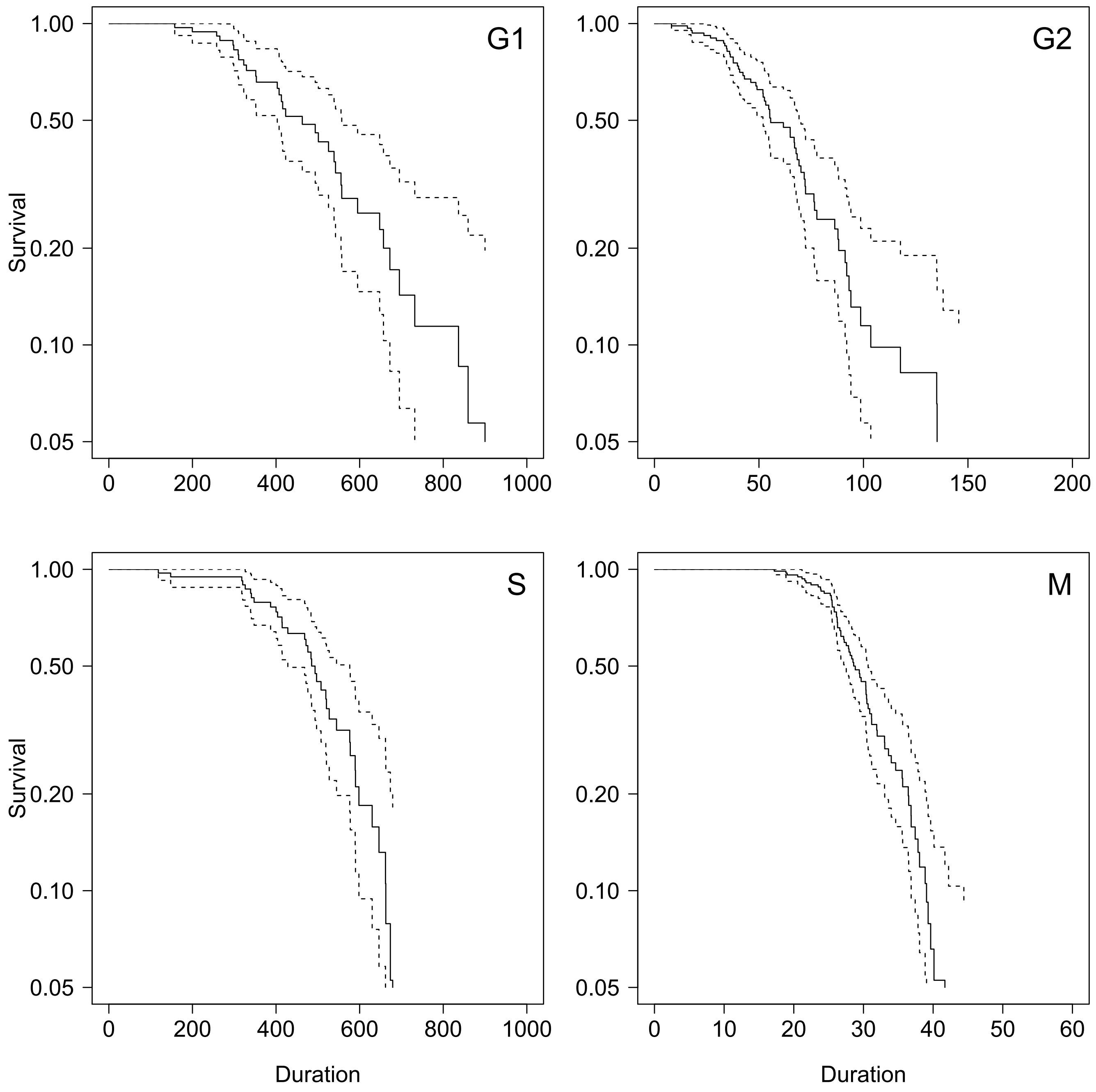

Figure 2: Phases Survival curves

##### 1.3.3 CDC25B$^{\Delta CDK}$

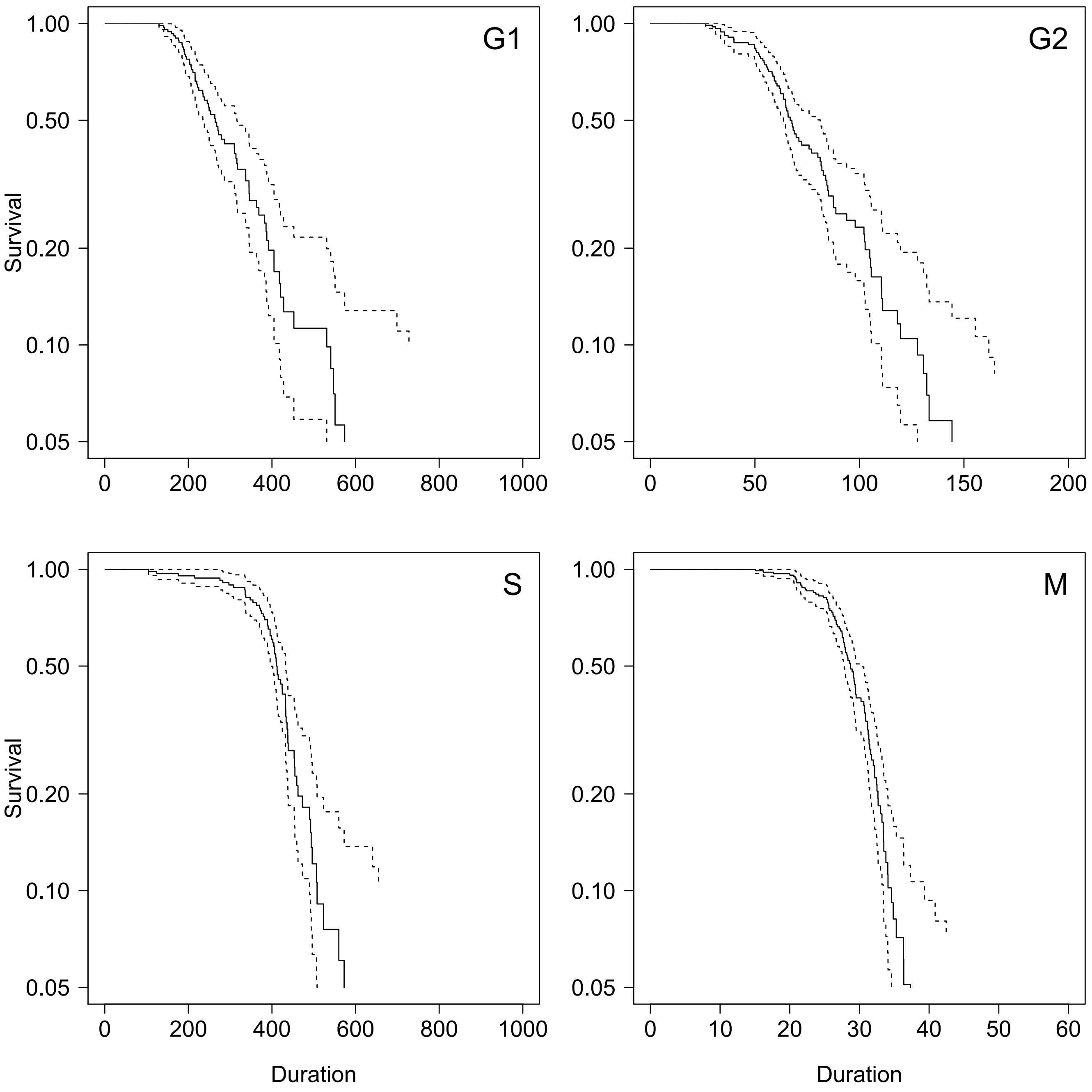

Figure 3: Phases Survival curves

#### 2 Subset with complete tracking of phases within the same cycle

In this section, we only use the subset of data with complete cell cycle measures.

##### 2.1 Graphical summaries

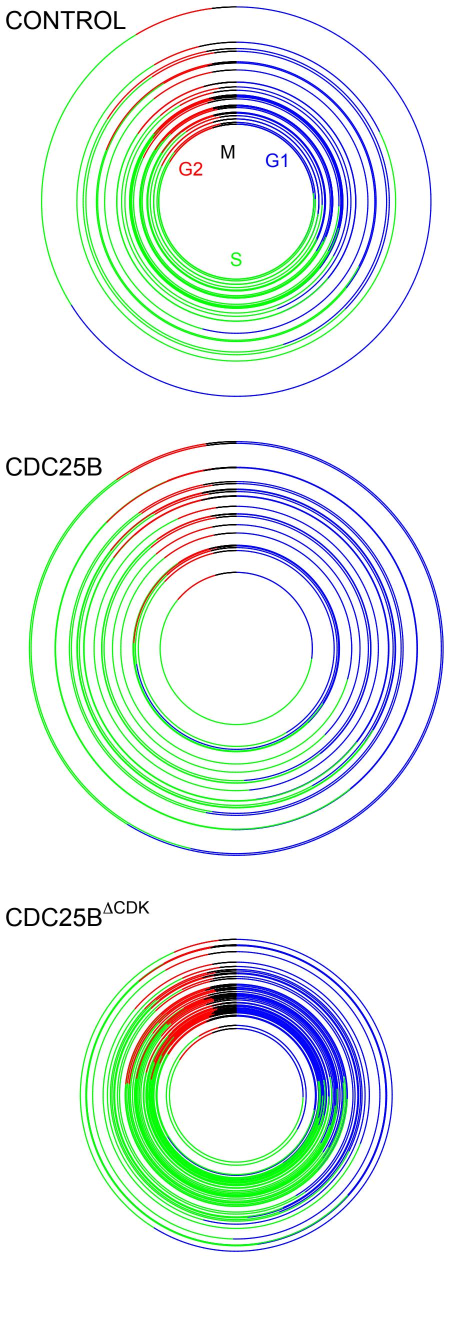

Figure 4: Graphical summary of data

#### 2.2 Variation Partitionning (Venn Diagrams)

The variation of cycle length is partionned in regards to phases to examine how much variation is explained by each phase.

See : Jari Oksanen, F. Guillaume Blanchet, Michael Friendly, Roeland Kindt, Pierre Legendre, Dan McGlinn, Peter R. Minchin, R. B. O'Hara, Gavin L. Simpson, Peter Solymos, M. Henry H. Stevens, Eduard Szoecs and Helene Wagner (2019). vegan: Community Ecology Package. R package version 2.5-6. https://CRAN.R-project.org/package=vegan

In Venn Diagrams, the number represent adjusted-Rsquare. For the sake of clarity, values below 0.05 are not reported.

In the three conditions, G1 phase duration explains the largest part of total length variations.

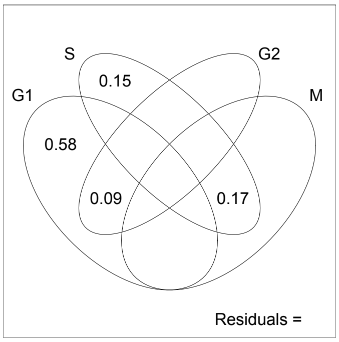

Figure 5: Venn Diagrams for CONTROL condition.

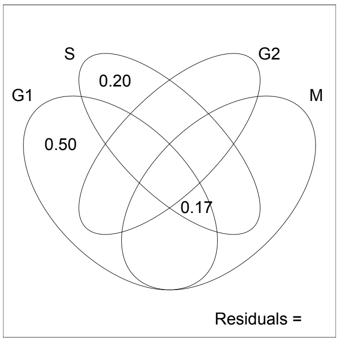

Figure 6: Venn Diagrams for CDC25B gain of fonction

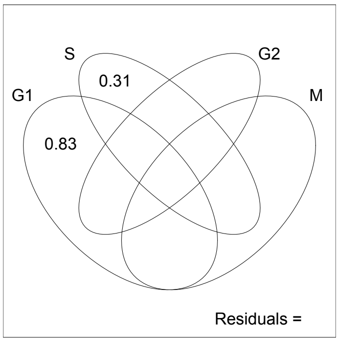

Figure 7: Venn Diagrams for DeltaCDK condition.

##### 2.3 Linear regression among phases durations

We test every pair of phases for linear regression. Tables below report $R^2$ values with their p-value, as well as correlation values with their associated p-value.

###### 2.3.1 CONTROL

| Phase | Phase | $R^2$ | p-val | Pearson | p-val | Kendall | p-val | Spearman | p-val | Signif |
|---|---|---|---|---|---|---|---|---|---|---|
| G1 | S | 0.00 | 0.90 | 0.02 | 0.90 | 0.17 | 0.18 | 0.23 | 0.20 | — |
| G1 | G2 | 0.09 | 0.08 | 0.30 | 0.08 | 0.22 | 0.07 | 0.33 | 0.06 | — |
| G1 | M | 0.01 | 0.57 | 0.10 | 0.57 | 0.09 | 0.45 | 0.17 | 0.35 | — |
| S | G2 | 0.00 | 0.80 | 0.05 | 0.80 | -0.00 | 0.99 | -0.01 | 0.94 | — |
| S | M | 0.49 | 0.00 | 0.70 | 0.00 | 0.33 | 0.01 | 0.47 | 0.01 | yes |
| G2 | M | 0.04 | 0.26 | 0.20 | 0.26 | 0.15 | 0.22 | 0.24 | 0.17 | — |
| G1 | SG2M | 0.01 | 0.58 | 0.10 | 0.58 | 0.25 | 0.04 | 0.35 | 0.05 | — |

###### 2.3.2 CDC25B

| Phase | Phase | $R^2$ | p-val | Pearson | p-val | Kendall | p-val | Spearman | p-val | Signif |
|---|---|---|---|---|---|---|---|---|---|---|
| G1 | S | 0.05 | 0.30 | 0.23 | 0.30 | 0.17 | 0.29 | 0.24 | 0.29 | — |
| G1 | G2 | 0.02 | 0.53 | 0.14 | 0.53 | 0.16 | 0.31 | 0.21 | 0.36 | — |
| G1 | M | 0.16 | 0.07 | 0.40 | 0.07 | 0.31 | 0.05 | 0.43 | 0.05 | — |
| S | G2 | 0.04 | 0.39 | 0.19 | 0.39 | 0.11 | 0.50 | 0.18 | 0.41 | — |
| S | M | 0.26 | 0.01 | 0.51 | 0.01 | 0.38 | 0.01 | 0.50 | 0.02 | yes |
| G2 | M | 0.05 | 0.31 | 0.23 | 0.31 | 0.19 | 0.22 | 0.26 | 0.23 | — |
| G1 | SG2M | 0.07 | 0.24 | 0.26 | 0.24 | 0.21 | 0.18 | 0.31 | 0.16 | — |

###### 2.3.3 CDC25B$^{\Delta CDK}$

| Phase | Phase | $R^2$ | p-val | Pearson | p-val | Kendall | p-val | Spearman | p-val | Signif |
|---|---|---|---|---|---|---|---|---|---|---|
| G1 | S | 0.01 | 0.56 | -0.09 | 0.56 | 0.13 | 0.22 | 0.18 | 0.24 | — |
| G1 | G2 | 0.03 | 0.28 | -0.17 | 0.28 | -0.10 | 0.33 | -0.19 | 0.21 | — |
| G1 | M | 0.01 | 0.48 | -0.11 | 0.48 | -0.05 | 0.65 | -0.09 | 0.58 | — |
| S | G2 | 0.00 | 0.98 | -0.00 | 0.98 | -0.00 | 0.98 | 0.01 | 0.97 | — |
| S | M | 0.03 | 0.27 | 0.17 | 0.27 | 0.18 | 0.09 | 0.26 | 0.09 | — |
| G2 | M | 0.01 | 0.59 | 0.08 | 0.59 | 0.01 | 0.93 | 0.03 | 0.84 | — |
| G1 | SG2M | 0.02 | 0.34 | -0.14 | 0.34 | 0.02 | 0.85 | 0.02 | 0.92 | — |

#### 2.4 Predicted survival curves for total duration under independence hypothesis.

Survival curves for total duration Tc will be used to comfort the observation that phase durations are not correlated.

For this, we can compare the observed distribution of total durations to a theoretical model established under the null hypothesis of no correlation among phases duration.

However, such a theoretical model can be obtained only for restrictive condition and is used here only for CTL condition.

An alternative is to build a theoretical distribution using the data (Monte Carlo permutations).

##### 2.4.1 Theoretical *null* model for total duration

Under the hypotheses that phase durations are independent from each other, and that exit time from each phase is a pure memoryless random process, we have a theoretical model for the statistical distribution of the total duration as a sum of four exponential distributions, each with its own parameter. Let denote $R_i$ the *exit rates* (the inverse of the average exit time), then the survival function of total exit rate would follow:

$$S(t) = 1 - \left[\prod_{i=1}^{4} R_i\right] \sum_{i=1}^{4} \frac{e^{-R_i t}}{\prod_{k=1,\neq i}^{n} (R_k - R_i)}$$

See : Bibinger, M. (2013). Notes on the sum and maximum of independent exponentially distributed random variables with different scale parameters. https://arxiv.org/abs/1307.3945 )

The expected distribution of Tc is obtained by adding the sum of the four minimal durations to this.

##### 2.4.2 Null model by Monte Carlo permutation

Inspired from Monte Carlo permutation tests, the expected distribution of Tc under phase duration independence hypothesis can be built from data by random picking among observed values (mixing among the observed cycles). If there were some kind of compensation between phases within cycles to ensure some regulation of Tc, then the distribution of total duration would be more homogeneous that the one obtained by such random permutation : the slope of survival curve for observed data would be steeper than survival curve of random sampling.

##### 2.4.3 Ordered permutations with full anti-correlation between G1 and S+G2+M

To illustrate how compensation of duration between G1 phase and the other phases would affect the survival curve and a more homogeneous series, we build the extreme case where the durations would be perfectly anti-correlated. To do this, we pair the G1 series sorted in ascending order with the S+G2+M series sorted in descending order.

#### 2.5 Survival curves for phase exit time in each condition.

Here we check whether exponential decay could apply.

##### 2.5.1 CONTROL

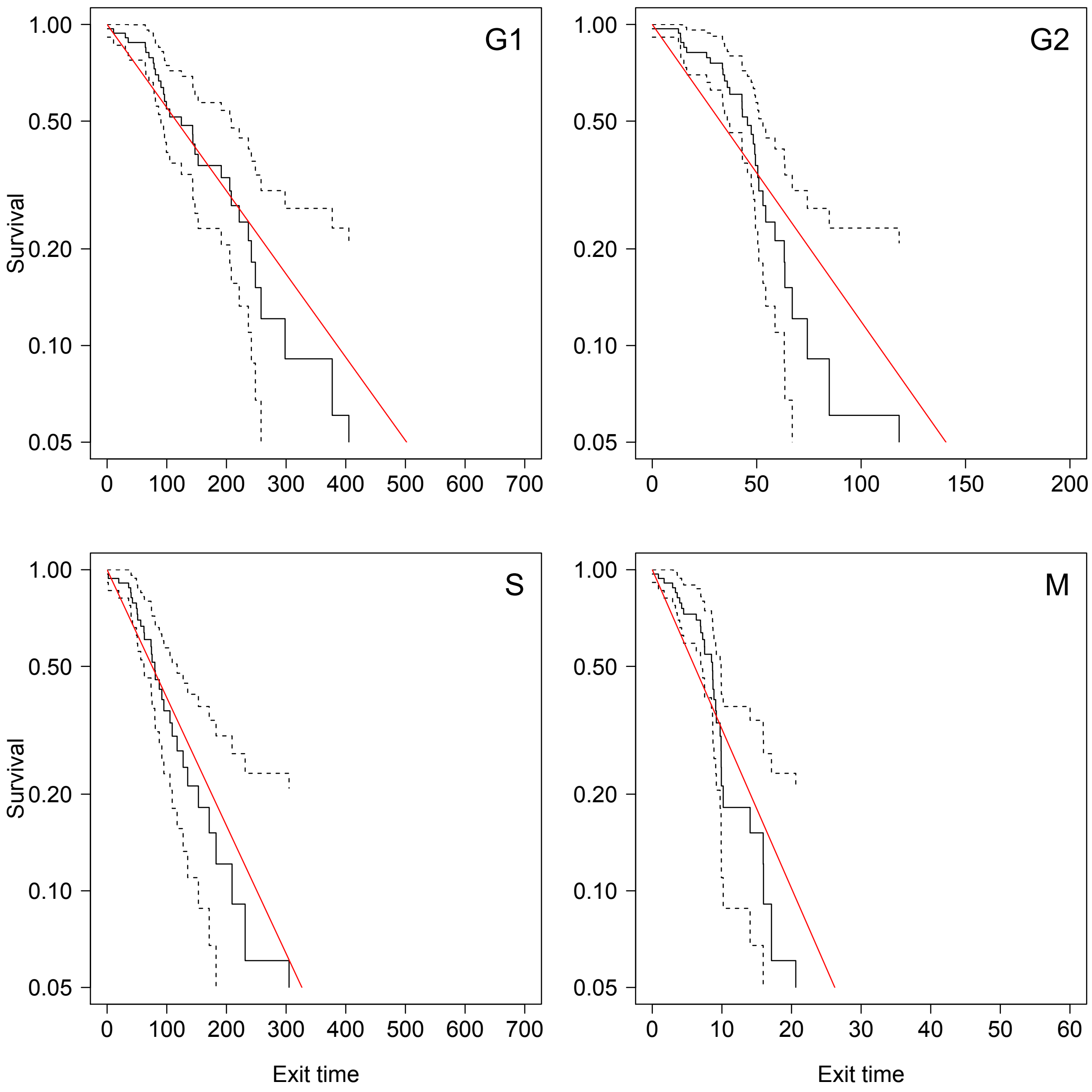

Figure 8: Phases Survival curves for exit time. If we substract the minimal duration, the exit time process seems compatible with an exponential decay (pure random memoryless process)

##### 2.5.2 CDC25B

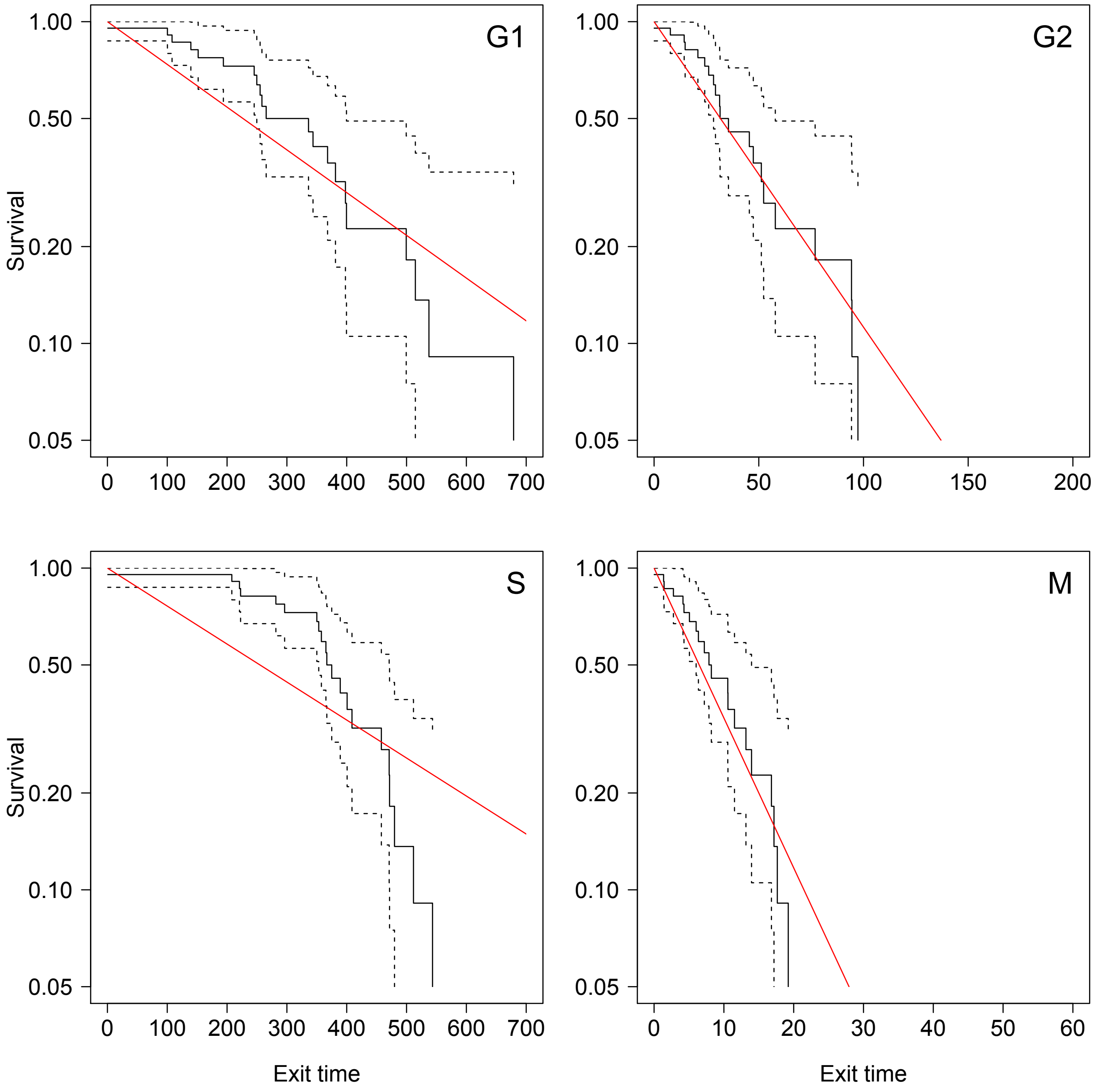

Figure 9: Phases Survival curves. The exit time process for S phase does not fit an exponential decay (accelerated process)

##### 2.5.3 CDC25B$^{\Delta CDK}$

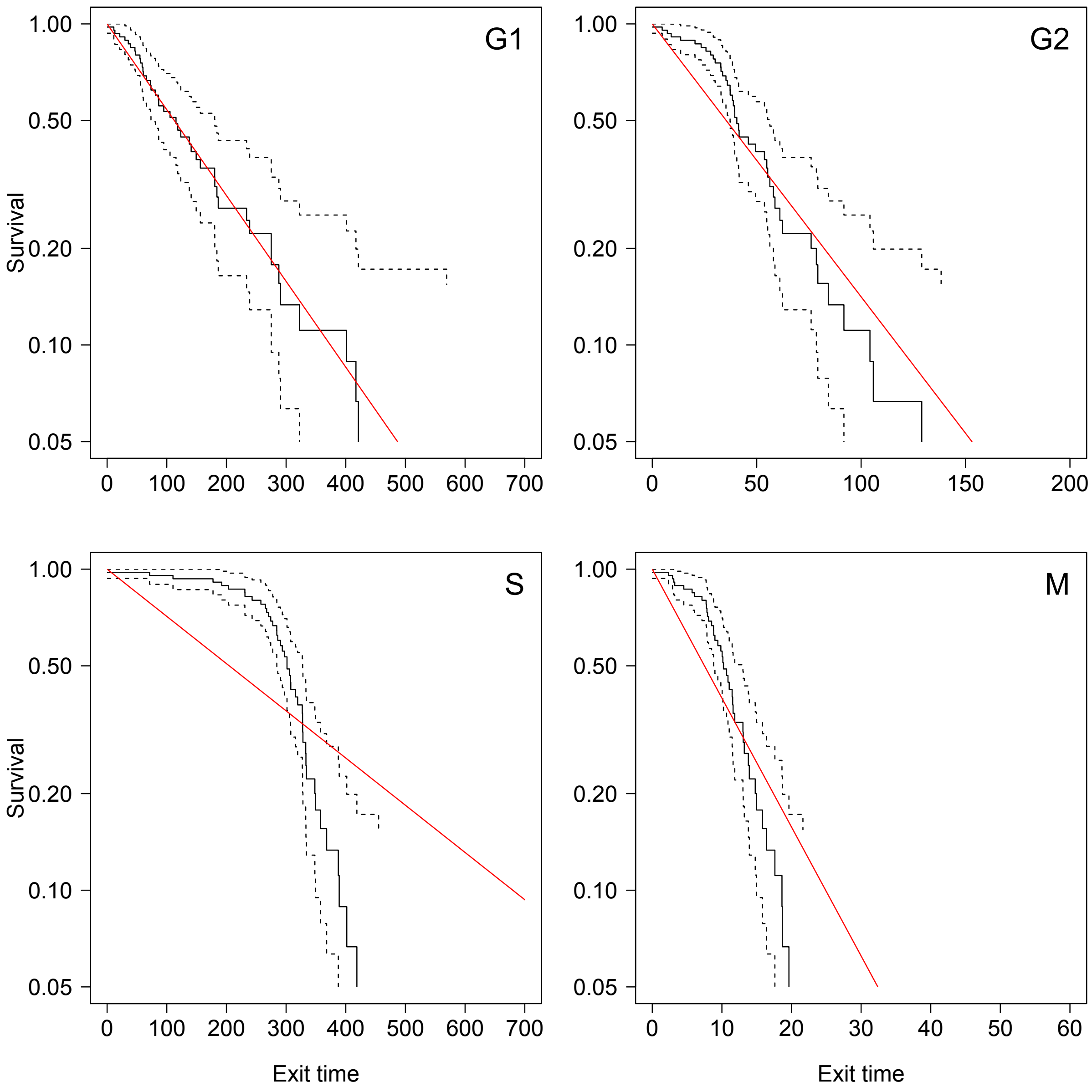

Figure 10: Phases Survival curves. The exit time process for S phase does not fit an exponential decay (accelerated process)

##### 2.6 Survival curves for Tc.

###### 2.6.1 CONTROL

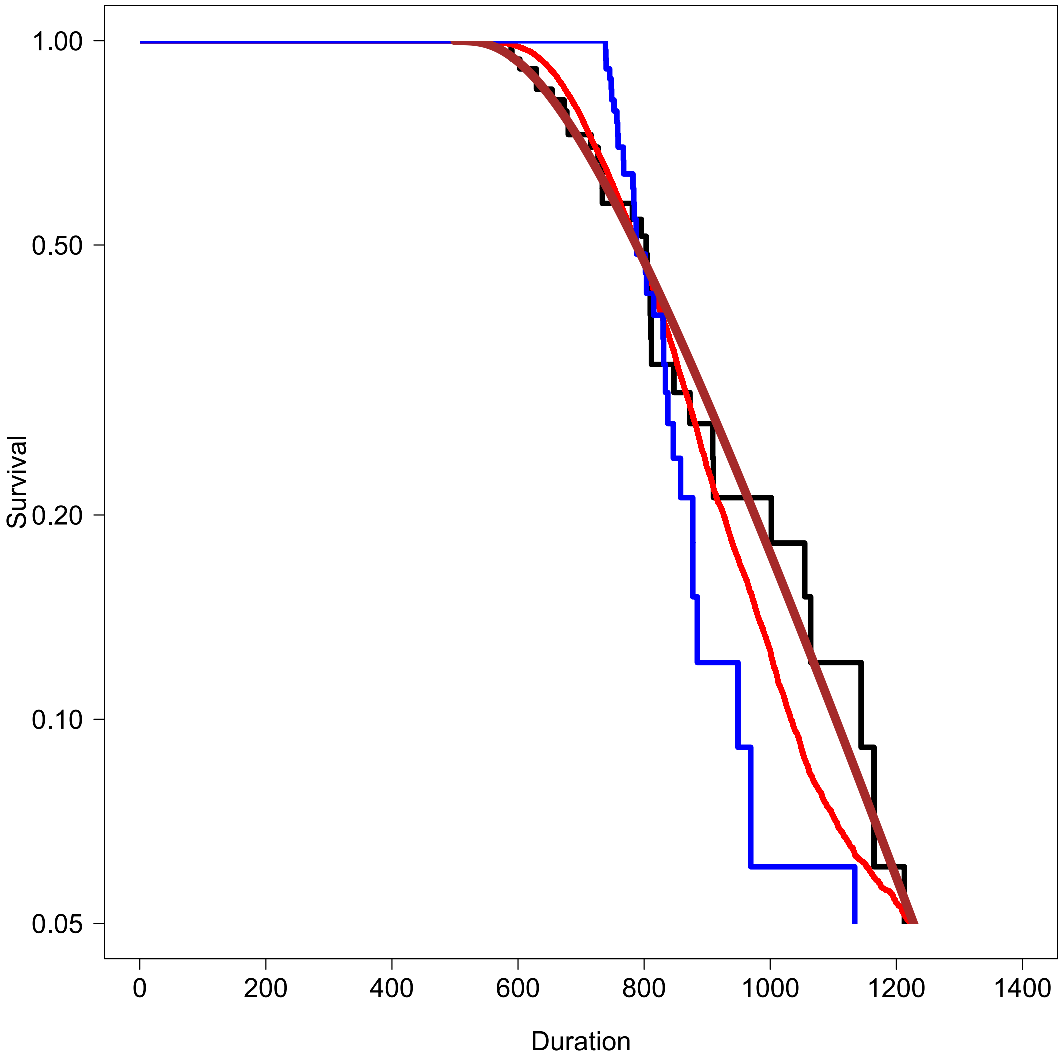

Figure 11: Tc Survival curve. Brown curve indicates the theoretical prediction of the null model of phase duration independence. Red curve indicates the survival obtained by Monte Carlo permutations of phase durations from the data set ($10^4$ samples). Blue curve indicates the survival for the anti-correlated pairing.

##### 2.6.2 CDC25B

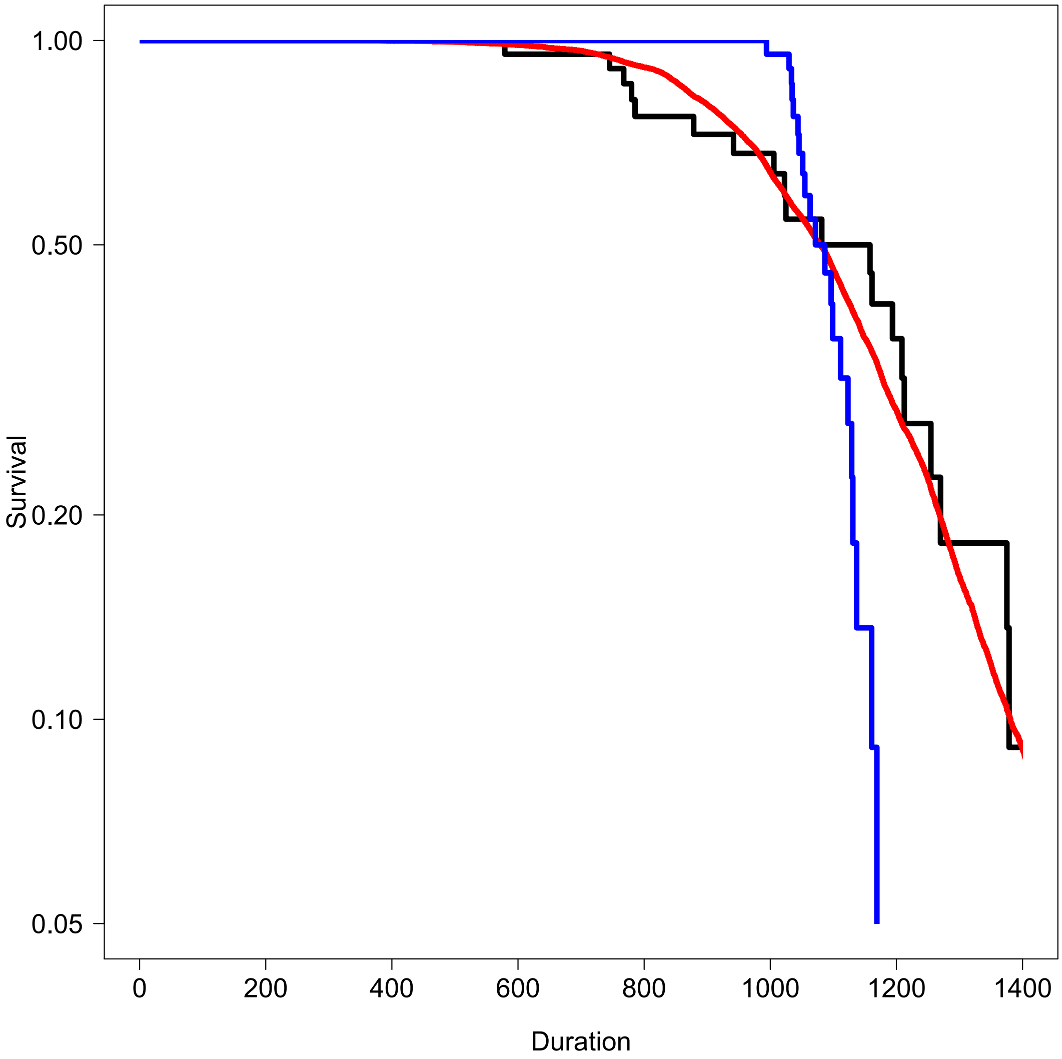

Figure 12: Tc Survival curve. Since the survival curves for phase durations do not match exponential decay, the theoretical null model does not apply and is not shown. Red curve indicates the survival of total lengths obtained by Monte Carlo permutations of phase durations from the data set. Blue curve indicates the survival for the anti-correlated pairing.

##### 2.6.3 CDC25B$^{\Delta CDK}$

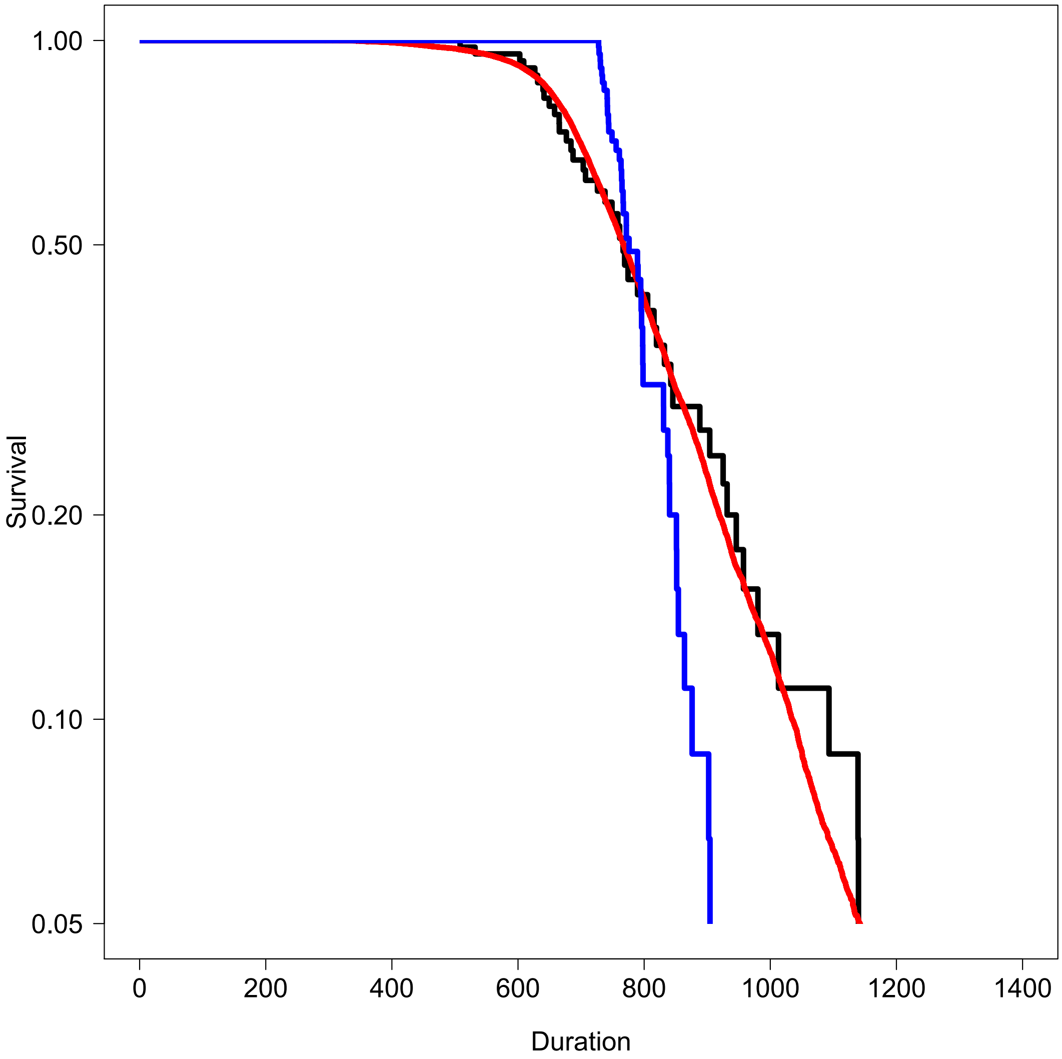

Figure 13: Tc Survival curve. Since the survival curves for phase durations do not match exponential decay, the theoretical null model does not apply and is not shown. Red curve indicates the survival of total lengths obtained by Monte Carlo permutations of phase durations from the data set. Blue curve indicates the survival for the anti-correlated pairing.

#### 3 Survival Analysis for the experimental treatment

For each condition, we report two graphical representations : a histogram representation (vertical dotted line reports the average of the distribution) and the corresponding survival curves.

The p-value of the survival test is given by the Score (logrank) test.

The output of Cox Proportional Hazard test is given for information to indicate the order of ratio between rates (e.g. exp(coef)=0.5 for CDC25B factor indicates that the exit rate is twice as slow as the baserate (CONTROL)).

For the sake of graphical clarity, confidence intervals are not reported.

For survival analysis of Tc, only cells with fully tracked cycles are used.

## 3.1 Tc

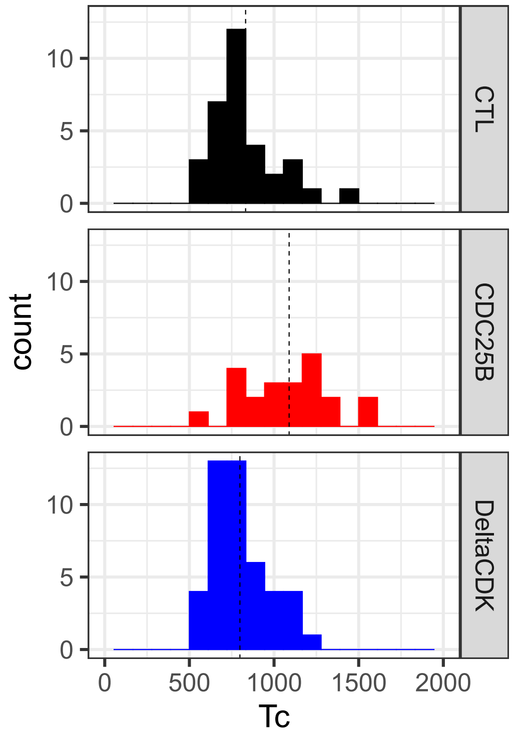

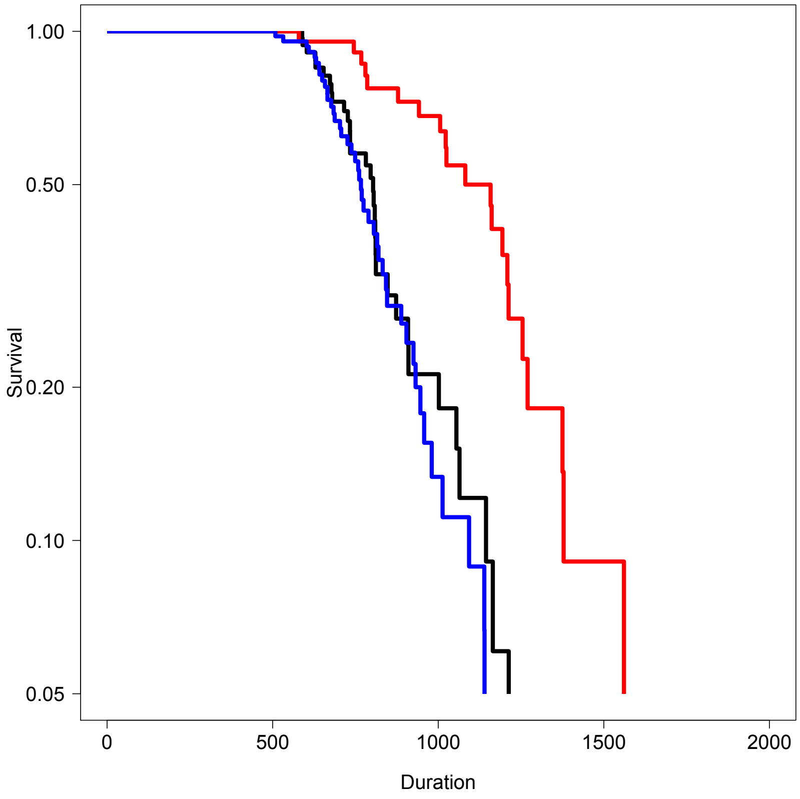

```
Call:
coxph(formula = Surv(dt$Tc, dt$sta) ~ dt$Condition)

  n= 100, number of events= 100
   (128 observations deleted due to missingness)

                          coef  exp(coef)  se(coef)      z  Pr(>|z|)
dt$ConditionCDC25B     -1.0393    0.3537    0.2932 -3.544 0.000394 ***
dt$ConditionDeltaCDK    0.2690    1.3086    0.2352  1.144 0.252695
---
Signif. codes:  0 '***' 0.001 '**' 0.01 '*' 0.05 '.' 0.1 ' ' 1

                     exp(coef) exp(-coef) lower .95 upper .95
dt$ConditionCDC25B      0.3537     2.8273    0.1991    0.6284
dt$ConditionDeltaCDK    1.3086     0.7641    0.8254    2.0749

Concordance= 0.615  (se = 0.031 )
Likelihood ratio test= 23.66  on 2 df,   p=7e-06
Wald test            = 20.13  on 2 df,   p=4e-05
Score (logrank) test = 21.74  on 2 df,   p=2e-05
```

## 3.2 G1

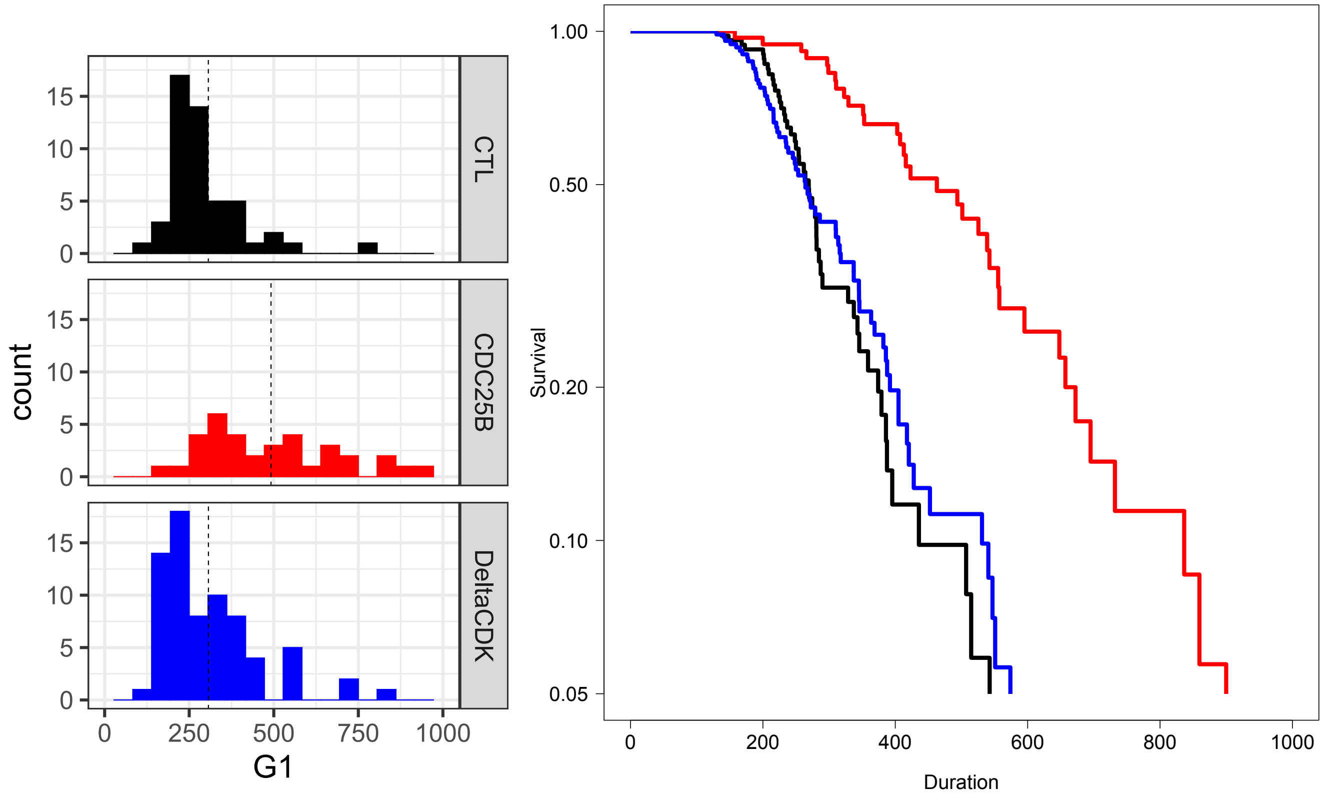

```
Call:
coxph(formula = Surv(dt$G1, dt$sta) ~ dt$Condition)

  n= 157, number of events= 157
   (71 observations deleted due to missingness)

                          coef exp(coef) se(coef)      z Pr(>|z|)
dt$ConditionCDC25B    -0.88337   0.41339  0.22647 -3.901  9.6e-05 ***
dt$ConditionDeltaCDK   0.01138   1.01145  0.18672  0.061    0.951
---
Signif. codes:  0 '***' 0.001 '**' 0.01 '*' 0.05 '.' 0.1 ' ' 1

                     exp(coef) exp(-coef) lower .95 upper .95
dt$ConditionCDC25B      0.4134     2.4190    0.2652    0.6444
dt$ConditionDeltaCDK    1.0114     0.9887    0.7015    1.4584

Concordance= 0.608  (se = 0.025 )
Likelihood ratio test= 22.92  on 2 df,   p=1e-05
Wald test            = 20.17  on 2 df,   p=4e-05
Score (logrank) test = 21.29  on 2 df,   p=2e-05
```

### 3.3 S

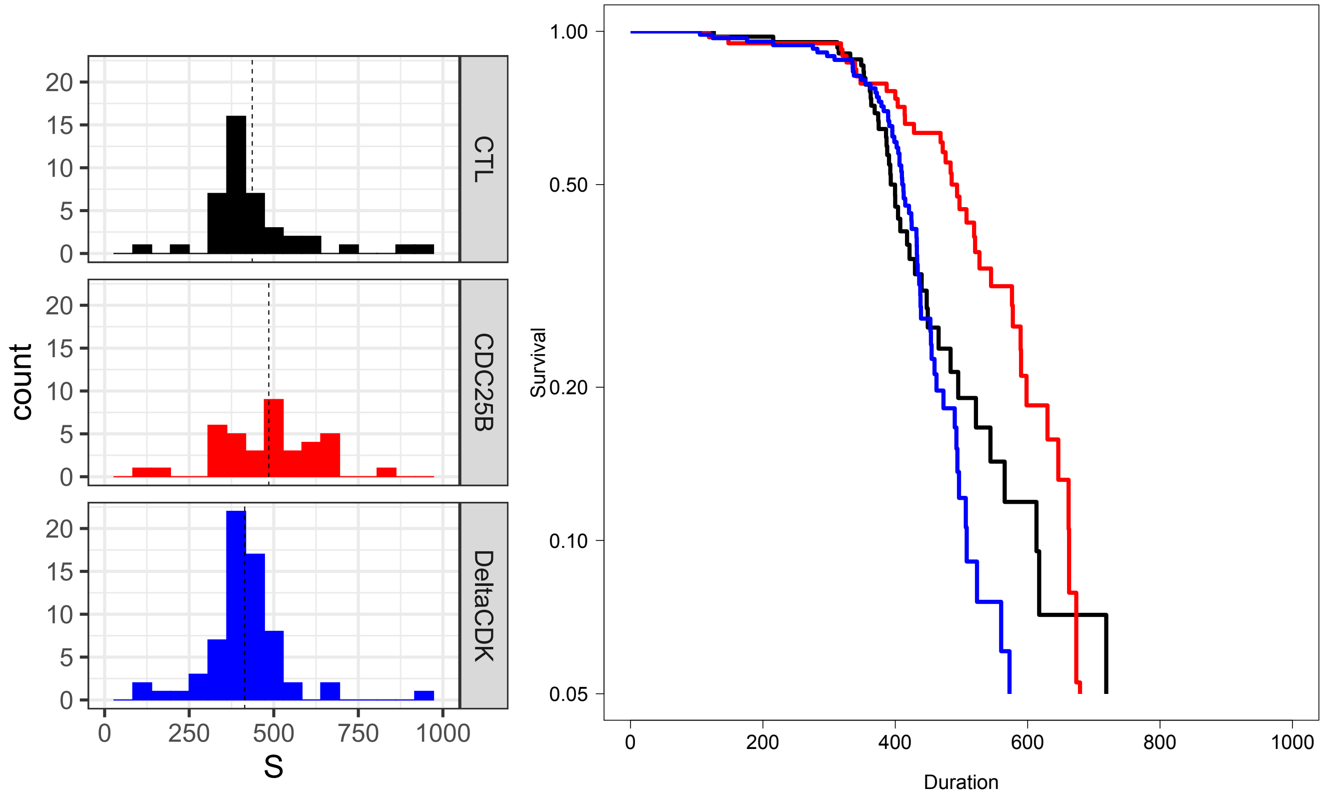

```
Call:
coxph(formula = Surv(dt$S, dt$sta) ~ dt$Condition)

  n= 146, number of events= 146
   (82 observations deleted due to missingness)

                         coef exp(coef) se(coef)      z Pr(>|z|)
dt$ConditionCDC25B   -0.4198    0.6572   0.2284 -1.838   0.0661 .
dt$ConditionDeltaCDK  0.1809    1.1983   0.2021  0.895   0.3706
---
Signif. codes:  0 ‘***’ 0.001 ‘**’ 0.01 ‘*’ 0.05 ‘.’ 0.1 ‘ ’ 1

                     exp(coef) exp(-coef) lower .95 upper .95
dt$ConditionCDC25B      0.6572     1.5217    0.4200     1.028
dt$ConditionDeltaCDK    1.1983     0.8345    0.8065     1.781

Concordance= 0.559  (se = 0.027 )
Likelihood ratio test= 8.69  on 2 df,   p=0.01
Wald test            = 8.25  on 2 df,   p=0.02
Score (logrank) test = 8.43  on 2 df,   p=0.01
```

## 3.4 G2

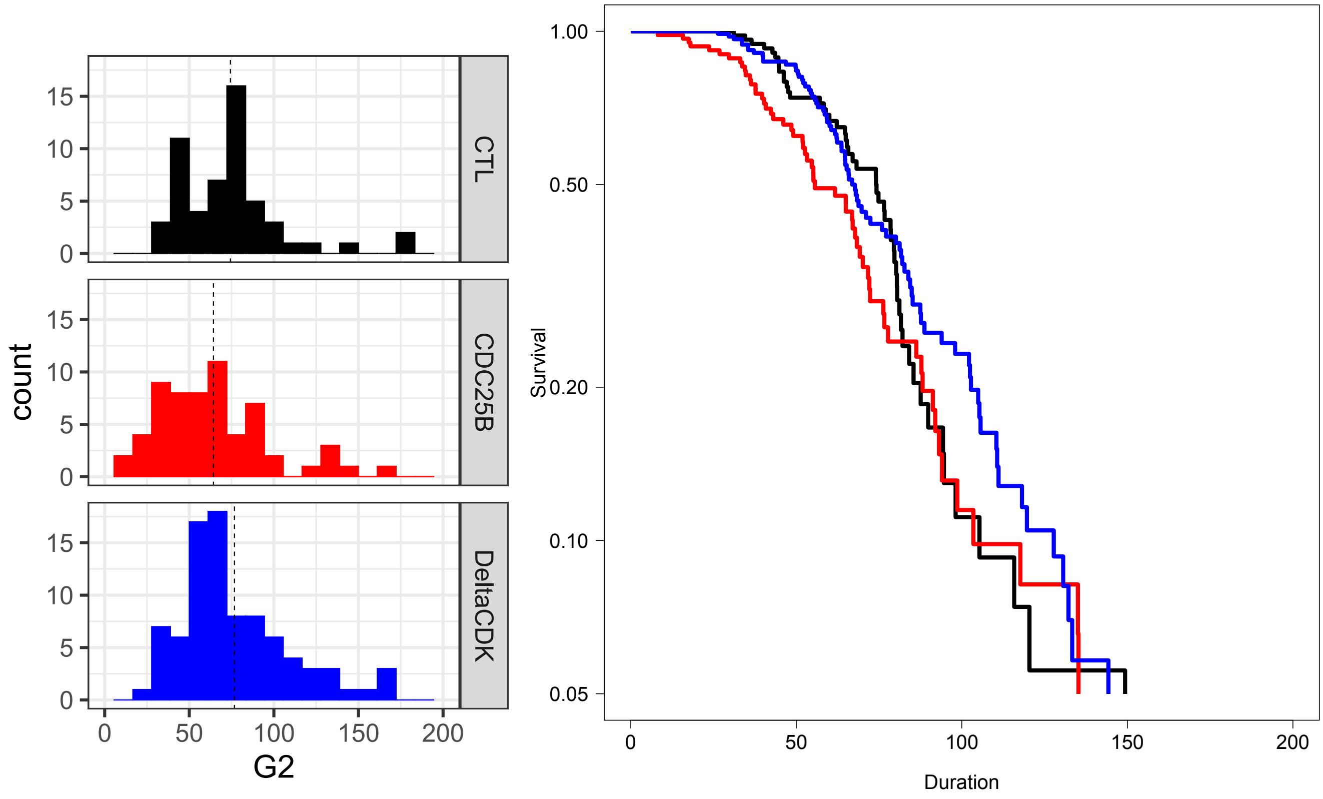

```
Call:
coxph(formula = Surv(dt$G2, dt$sta) ~ dt$Condition)

  n= 201, number of events= 201
   (27 observations deleted due to missingness)

                         coef exp(coef) se(coef)      z Pr(>|z|)
dt$ConditionCDC25B    0.26603   1.30478  0.18989  1.401    0.161
dt$ConditionDeltaCDK -0.02233   0.97792  0.17675 -0.126    0.899

                     exp(coef) exp(-coef) lower .95 upper .95
dt$ConditionCDC25B      1.3048     0.7664    0.8993     1.893
dt$ConditionDeltaCDK    0.9779     1.0226    0.6916     1.383

Concordance= 0.55  (se = 0.023 )
Likelihood ratio test= 3.19  on 2 df,   p=0.2
Wald test            = 3.31  on 2 df,   p=0.2
Score (logrank) test = 3.33  on 2 df,   p=0.2
```

## 3.5 M

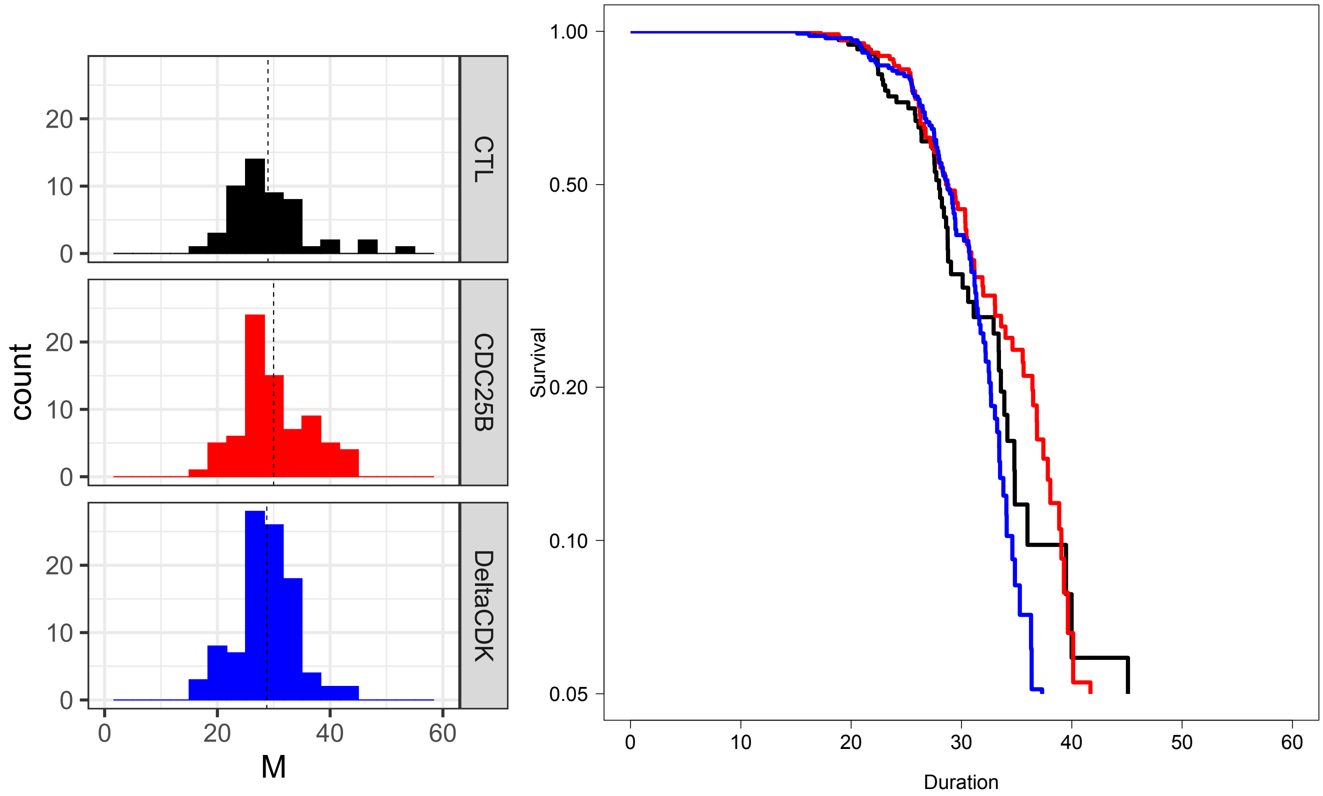

```
Call:
coxph(formula = Surv(dt$M, dt$sta) ~ dt$Condition)

  n= 225, number of events= 225
   (3 observations deleted due to missingness)

                         coef exp(coef) se(coef)      z Pr(>|z|)
dt$ConditionCDC25B   -0.07976   0.92334  0.18570 -0.430    0.668
dt$ConditionDeltaCDK  0.13998   1.15025  0.17888  0.783    0.434

                     exp(coef) exp(-coef) lower .95 upper .95
dt$ConditionCDC25B      0.9233     1.0830    0.6416     1.329
dt$ConditionDeltaCDK    1.1502     0.8694    0.8101     1.633

Concordance= 0.513  (se = 0.021 )
Likelihood ratio test= 2.07  on 2 df,   p=0.4
Wald test            = 2.07  on 2 df,   p=0.4
Score (logrank) test = 2.08  on 2 df,   p=0.4
```

### Figure sup 3

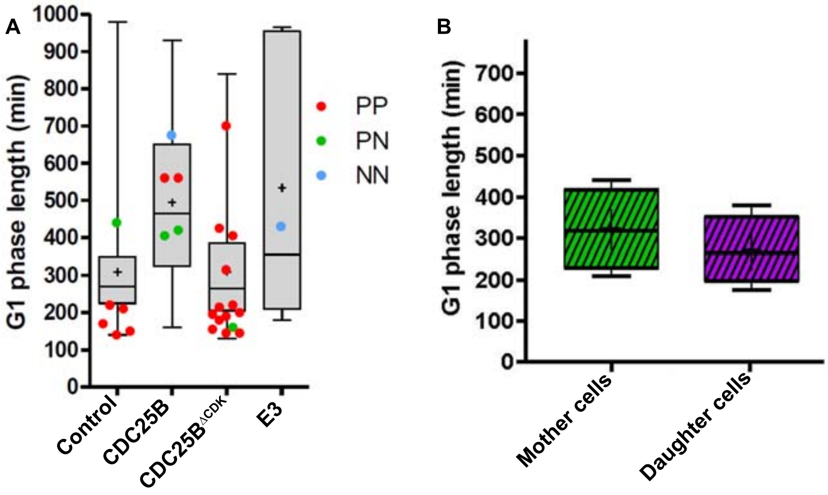
A
G1 phase length (min)
1000
900
800
700
600
500
400
300
200
100
0
Control
CDC25B
CDC25B$^{\Delta CDK}$
E3
PP
PN
NN
B
G1 phase length (min)
700
600
500
400
300
200
100
0
Mother cells
Daughter cells

Figure sup 4

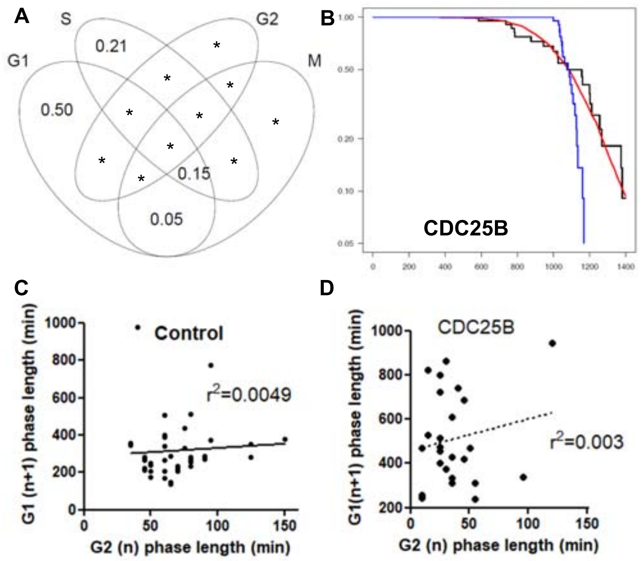
A
S
G2
G1
M
0.21
0.50
0.15
0.05
B
CDC25B
C
Control
$r^2$=0.0049
G1 (n+1) phase length (min)
G2 (n) phase length (min)
D
CDC25B
$r^2$=0.003
G1(n+1) phase length (min)
G2 (n) phase length (min)
